# Antibody-drug conjugates to treat bacterial biofilms

**DOI:** 10.1101/2023.01.16.524127

**Authors:** Anne Tvilum, Mikkel I. Johansen, Lærke N. Glud, Diana M. Ivarsen, Amanda B. Khamas, Sheiliza Carmali, Snehit Satish Mhatre, Ane B. Søgaard, Emma Faddy, Lisanne de Vor, Suzan H.M. Rooijakkers, Lars Østergaard, Nis P. Jørgensen, Rikke L. Meyer, Alexander N. Zelikin

**Affiliations:** Department of Chemistry, Aarhus University, Aarhus C 8000, Denmark; Department of Clinical Medicine, Aarhus University, Aarhus N 8200, Denmark; Department of Infectious Diseases, Aarhus University Hospital, Aarhus N 8200, Denmark; Interdisciplinary Nanoscience Centre (iNANO), Aarhus University, Aarhus C 8000, Denmark; Department of Medical Microbiology, University Medical Center Utrecht, Utrecht, The Netherlands; Department of Biology, Aarhus University, Aarhus C 8000, Denmark

## Abstract

Implant-associated infections remain a grand unmet medical need because they involve biofilms that protect bacteria from the immune system and harbour antibiotic-tolerant persister cells. There is an urgent need for new biofilm-targeting therapies with antimicrobials, to treat these infections via a non-surgical way. In this work, we address this urgent medical need and engineer antibody-drug conjugates (ADC) that kill bacteria in suspension and in biofilms, *in vitro* and *in vivo*. The ADC contains an anti-neoplastic drug mitomycin C, which is also a potent antimicrobial against biofilms. While most ADCs are clinically validated as anti-cancer therapeutics where the drug is released after internalisation of the ADC in the target cell, the ADCs designed herein release the conjugated drug without cell entry. This is achieved with a novel mechanism of drug, which likely involves an interaction of ADC with thiols on the bacterial cell surface. ADC targeted towards bacteria were superior by the afforded antimicrobial effects compared to the non-specific counterpart, in suspension and within biofilms, *in vitro* and *in vivo*. An implant-associated murine osteomyelitis model was then used to demonstrate the ability of the antibody to reach the infection, and the superior antimicrobial efficacy compared to standard antibiotic treatment *in vivo*. Our results illustrate the development of ADCs into a new area of application with a significant translational potential.

## Introduction

Bacterial colonization of implanted biomaterials leads to infections that are a serious complication with a high socio-economic and healthcare burden. Despite advances in surgery, infection remains a risk, with incidence rates of 1-2% for prosthetic knees and hips,^1^ 1-5% for prosthetic vascular grafts^2^ and up to 8.5% for spinal implants^3^. Post-operative prophylaxis often has little to no effect on the implant-associated infections, ^4^ and surgical intervention is often required to cure the patient. ^5^ Patients ineligible for surgery are faced with either amputation of limbs or lifelong suppressive antibiotic therapy, which is also associated with significant morbidity.

The resilience of implant-associated infections is linked to the formation of bacterial biofilms on and around the implant. ^5,6^ Bacteria in biofilms are embedded in a shared extracellular matrix, which offers protection from the immune system. Within the biofilm, slow-growing or dormant sub-populations emerge, and these populations are called “persister cells” because they survive extremely high concentrations of all the antibiotics in current clinical use. ^7^ Treatment of implant-associated infections therefore remains a major healthcare challenge and requires a novel treatment paradigm.^8^

Novel therapies developed to specifically tackle biofilm infections often rely on one of three strategies: 1) discovery of new antibiotics ^9^ or 2) delivery of a high local dose of current antibiotics, ^10^ or 3) prodrug therapy where an inactive drug is circulating in the body and only activated at site of infection ^11 12^ Another approach to tackle bacterial infections is gaining momentum, based on the use of phages as a nature-derived, bacteria-specific treatment.^13^ In this study we take advantage of drug repurposing and prodrug therapy to deliver an anti-neoplastic drug that is highly effective against biofilms and will benefit from a prodrug therapy approach to minimize side effects. Specifically, we develop an antibody-drug conjugate (ADC) of mitomycin C for the treatment for implant-associated biofilm infections caused by *Staphylococcus aureus*, which is the most common culprit in prosthetic joint infections.^6^

The motivation for this endeavour lies in that ADCs are among the most successful tools of biomedicine, specifically for targeted drug delivery in cancer therapy.^14,15^ Within these prodrugs, the antibody arm is responsible for the association with a cognate cell surface ligand, judiciously chosen as a marker of a disease. In the majority of successful designs of ADCs, the linker between the antibody and the drug is designed to be stable during prodrug circulation in the blood and to be degraded upon cell entry.^16^ This measure ensures the highest specificity of action for an ADC against the nominated target. From the standpoint of chemistry, this is achieved using linkers that are stable in the oxidative environment of blood and neutral pH. Intracellular drug release is then achieved in response to acidification during the endosomal-lysosomal trafficking of the ADC, using pH sensitive linkers between the protein and the drug. Significantly higher intracellular concentration of the thiol-containing tripeptide glutathione (GSH) over its extracellular content makes disulfide linkages highly successful in the design of ADCs, in which case drug release occurs in the cell cytosol via thiol-disulfide exchange. Finally, arguably the most successful linker methodology relies on peptide sequences that are degraded by the intracellular proteases such as cathepsin B. ^16^ ADCs represent a highly successful paradigm with at least fourteen products approved to market, ^14,15,17^ all of which are developed for intracellular drug delivery and the treatment of cancer. Arguably the most prominent example of an ADC for the treatment of bacterial diseases (also termed antibody-antibiotic conjugates, AAC) is the one whereby the antibody binds a bacterium and through opsonization facilitates the uptake of the pathogen by the immune cells. ^18,19^ Here, a rifamycin analogue was conjugated to this antibody using a cathepsin-sensitive linker such that drug release was mediated by mammalian proteases, within the immune cells. ^18,19^ Apart from this successful case, examples of ADCs developed for the treatment of bacteria are few. ^20^ In large part, this is because bacteria do not perform receptor-mediated endocytosis.

We hypothesized that ADC can be developed into a powerful tool in the fight against bacterial pathogens, if drug release was engineered as an extracellular process. Extracellular drug release in the near-vicinity of the targeted cell is a highly promising concept with applications in cancer treatment. ^21–23^ Triggered extracellular drug release can be engineered with the knowledge of the enzymatic fingerprint of a disease, e.g. cancer, via the enzyme-prodrug therapy.^24^ ADCs have also been engineered to release the drug at the cell surface, in response to the enzymes attached exofacially to the cell surface.^22^ Key requirement with regards to the choice of the drug for these applications is that this therapeutic molecule has to be well cell permeable.

We hypothesized that this approach can be particularly well suited in the design of ADCs towards treatment of the implant associated bacterial infections. An antibody can be selected to anchor the prodrug to the surface of the bacteria within the biofilm before triggering release at the bacterial cell surface, using an external small molecule or via an interaction with the specific components of the bacterial cell surface and/or biofilm. This approach potentially opens the door to the use the most potent antimicrobials that are effective against biofilm infections, but have not been implemented in the clinic due to their side effects. Local drug release within the biofilm will minimize the systemic concentration of active drug while maximising its therapeutic impact.

Small molecule-triggered therapeutic activity has been applied to on-demand activation of the chimeric antigen receptor T cells (CAR T),)^25 26^ to dissolve implanted hydrogels and thus to release the second dose in a prime-boost vaccination approach,^27^ as well as for drug release from the “click-to-release” linkers.^21^ These elegant techniques rely on safe molecules, possibly marketed as therapeutics, to act as a switch in a molecularly programmed gate. We hypothesized that a marketed mucolytic agent, N-acetyl cysteine, can act as such a switch to trigger drug release from the ADCs that in their structure feature a disulfide-linkage between the antibody and the drug.

From a different perspective, we also considered that cell surfaces, mammalian ^28–30^ and bacterial ^31–33^, are often characterized with abundant accessible thiols. For mammalian cells, these thiols have been used to initiate drug release and to achieve cell entry, through conjugation to the cell surface thiols via thiol-disulfide exchange and subsequent internalization of the cell-bound cargo. ^29,34^ To the best of our knowledge, this technique has not been applied to bacterial cells, though the presence of thiols on the surface of bacterial outer membrane (for the Gram-negative bacteria) or the cell wall (for Gram-positive pathogens) has been experimentally confirmed in a number of studies. ^31–33^ It therefore seemed highly plausible that these surface thiols may be suited to initiate drug release via thiol-disulfide exchange.

The above presented design considerations dictated the final composition of ADCs engineered in this work. We used commercially available antibodies against *S. aureus* and synthesized ADCs with a disulfide linkage between the antibody and the drug mitomycin C (Figure 1A). Mitomycin C was chosen due to its outstanding antimicrobial activity against biofilms and the persister cells within.^35^ Towards the overall goal, we i) established the mitomycin C containing ADC and validated drug release from these prodrugs in response to N-acetyl cysteine (NAC) and a panel of other biologically relevant thiols; ii) visualized specific interaction of *S. aureus-specific* ADCs with planktonic bacteria and biofilms, iii) quantified the antibacterial efficacy of ADCs against planktonic bacteria and biofilms, and iv) demonstrated *in vivo* therapeutic effects of the ADCs in a murine implant-associated osteomyelitis model. Unexpectedly, we established that drug release is independent of NAC in the presence of bacteria, and we attribute the bacteria-triggered drug release to interaction of the ADCs with thiols on the bacterial cell surface. The importance of our data is in that we reengineer the highly successful therapeutic tool, ADC, to the previously un-explored area of use, and establish a novel modality of treatment against condition with a high medical need. We anticipate that these data will have a strong academic impact and believe that the ADC have a promising translational perspective.

**Figure 1.**
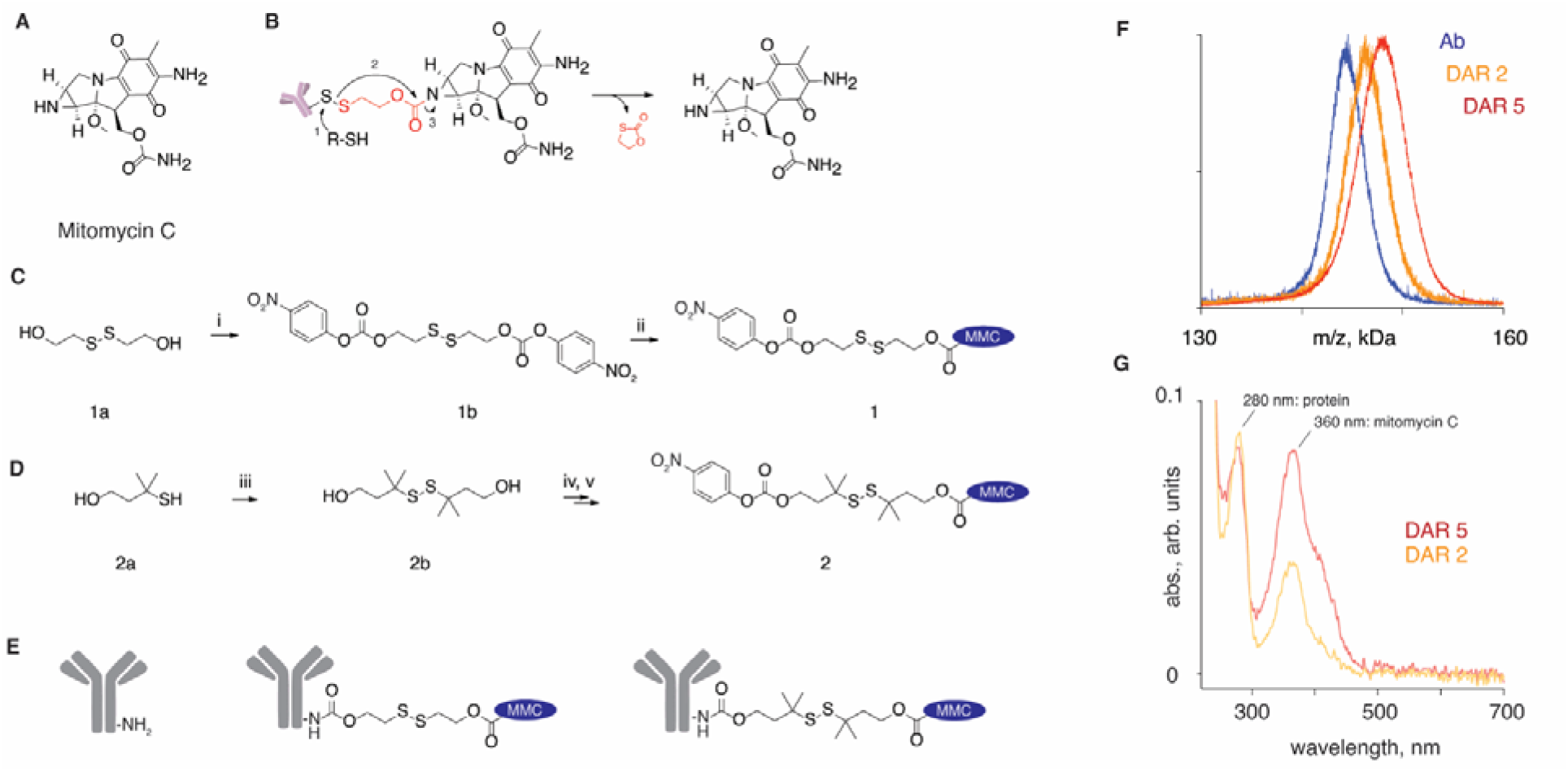
(A) Chemical formula of mitomycin C; (B) schematic illustration of the use of self-immolative linker to achieve release of the thiol-free drug, mitomycin, from its prodrugs (e.g. antibody-drug conjugates) triggered via thiol-disulfide exchange; (C-E) schematic illustration of syntheses of the amine-reactive, mitomycin C containing linker-drug conjugates **1** and **2**, for a single step conjugation to antibodies. Experimental conditions: i) **1a** (1 equiv.), 4-Nitrophenyl chloroformate (2.2 equiv.), TEA (4 equiv.), CH_2_Cl_2_, 0 ^o^C to r.t., 2 hours, 61 %; ii) **1b** (2.4 equiv.), Mitomycin C (1 equiv.), TEA (1.5 equiv.), HOBt (6.7 equiv.), DMF, 24 hours, 57 %; iii) **2a** (1 equiv.), CuCl_2_ (5 equiv.), CH_2_Cl_2_:EtOAc (1:1), HCl (0.5 M), 1.5 hours, r.t., 73 %; iv) **2b** (1 equiv.), 4-Nitrophenyl chloroformate (2.8 equiv.), TEA (2.5 equiv.), CH_2_Cl_2_, 3 hours, r.t., 69 %; v) **2c** (1 equiv.), Mitomycin C (1 equiv.), TEA (1.1 equiv.), HOBt (1.5 equiv.), CH_2_Cl_2_, 0 °C to r.t. over 10 min., r.t. for 24 hours, 25 %; conjugation to antibodies: Ab (c_protein_ > 3 mg/ml), **1** or **2** (55 equiv.), 10 mM PBS, pH 7.4, 2 hours; (F,G) Characterization of the ADC with drug-to-antibody (DAR) ratio of 2 and 5 using MALDI (**F**) and UV-vis spectroscopy (**G**).

## Results

### Design of ADC

Disulfide chemistry is among the most investigated linker technologies in drug delivery. The one limitation commonly encountered in these endeavours is that a disulfide is a bond between two sulfur atoms, and only few marketed drugs feature thiols in their structure. A successful route to overcome this limitation and apply disulfide chemistry to a wide range of drugs is the use of “self-immolative linkers”. ^36^ These linkers are designed to undergo fast decomposition upon a scission of the disulfide bond, most commonly via intramolecular cyclization, to afford a traceless release of the drug from its conjugate in its pristine form (Figure 1B). Mercaptoethanol dimer (compound **1a,** Figure 1C) is a convenient, readily available starting material for these syntheses. In our work, **1a** was converted into a symmetrical homobifunctional amine-reactive carbonate, and subsequently reacted with mitomycin C to afford a drug-linker conjugate (**1**) for a single-step conjugation to an antibody. Literature survey reveals that stability of disulfides against decomposition in blood significantly improves when carbon atom(s) adjacent to sulfurs are substituted with e.g. methyl groups, which create steric shields to non-specific exchange between this disulfide and cysteine thiols on albumin and other serum proteins. Learning from this, linker **2** was designed, starting with 3-methyl-3-sulfanylbutan-1-ol (compound **2a,** Figure 1D) which was oxidized using CuCl_2_ into the sterically hindered disulfide (**2b**). This was reacted with 4-nitrophenyl chloroformate to afford the homo-bifunctional carbonate **2b** and then with mitomycin C to obtain the final drug-linker conjugate **2**.

Compounds **1** and **2** are amine-reactive and were used to conjugate to the antibodies in their aqueous solutions (Figure 1E). ADC were purified via gel filtration and characterized for composition using MALDI, which revealed that protein conjugates had molar mass higher than the pristine antibody and allowed to calculate the drug-to-antibody ratio (DAR) for the conjugates (results for ADC based on linker **1**, DAR= 2 and 5 are shown as examples in Figure 1F. UV-vis spectra of the ADC also revealed signature absorbance for both, the protein and the conjugated drug, with absorbance for the latter expectedly increasing with DAR (Figure 1G).

### Drug release

To investigate drug release, ADC were treated with a natural reducing agent, a thiol-containing tripeptide glutathione (GSH, 5 mM). The resulting mixture was separated by size via spin filtration and then the filtrate and the concentrate were independently analysed via UV-vis spectroscopy (Figure 2A). The concentrate (high molar mass) solute volume revealed the presence of the protein with minor residual content of the drug, while the filtrate (low molar mass) exhibited the UV-vis signature of mitomycin C without the protein. These data validate the success in the design of ADC that release their conjugated cargo via thiol-disulfide exchange.

**Figure 2.**
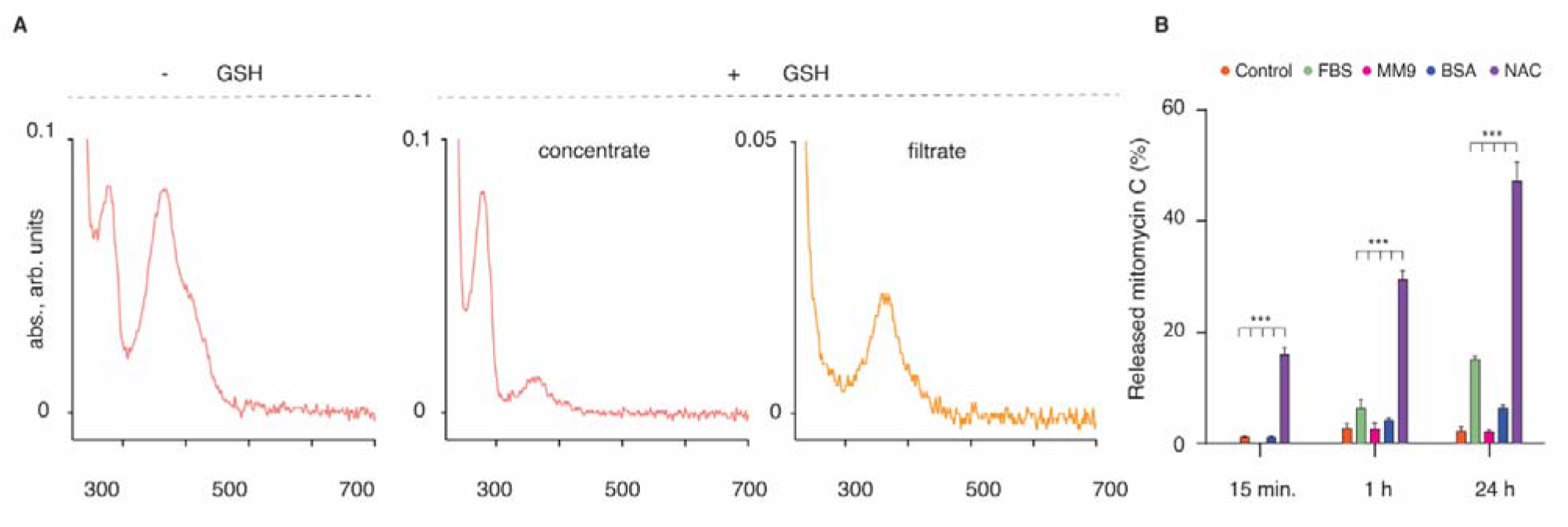
(A) Qualitative investigation of drug release from ADC using UV-vis spectroscopy showing the spectrum of the ADC (-GSH) and the spectra after the treatment with GSH and separation of the mixture by gel filtration, separately for the concentrate (the protein) and the filtrate (the drug); (B) drug release from ADC in PBS containing NAC (5 mM), bovine serum albumin (BSA, 0.76 μM), bacterial culture medium MM9, or FBS (10% in RPM1-1640 cell culture medium), over 24 h of incubation; control = PBS only; results are based on three independent experiments and shown as mean±st.dev.; statistical significance determined using two-way ANOVA analysis; for brevity, only selected significance values are presented, ***: *p* < 0,001.

Quantitative analysis of drug release from ADC was performed using High-Performance Liquid Chromatography (HPLC) (Figure 2B). To this end, the ADC was incubated in phosphate buffered saline (PBS) in the presence of NAC (5 mM), albumin (0.76 μM), or fetal bovine serum (10 % in RPM1-1640 cell culture medium), as well as in MM9 bacterial cell culture medium which was later to be used in antimicrobial testing. The common biochemical reducing agent dithiothreitol (DTT, 5 mM) was used as a positive control to achieve complete drug release from ADC. During the initial 15 minutes of incubation, ADC based on linker **1** exhibited negligible drug release in all conditions except for a treatment with NAC. With longer incubation, drug release became noticeable from ADC incubated in the presence of FBS, reaching approximately 20% after 24 h. Nevertheless, release triggered by NAC was substantially higher at all the time points. For the ADC designed using linker **2**, no drug was released upon incubation of ADC for 24 h in FBS or in PBS with BSA or NAC (data not shown). The only reducing agent that released mitomycin C was DTT, which has limited if any potential of use *in vivo*. Based on these results, in all subsequent experiments, we used ADC based on linker **1.**

### ADCs associate with the surface of *S. aureus* and retain immune-activating properties

Next, we aimed to validate antibody binding to bacteria. The optimal antibody for a design of ADC to deliver drugs to bacteria should bind to bacterial cells when growing planktonically and as biofilm, despite the differences in the expression of surface proteins in these two growth modes. To investigate this, we fluorescently labelled commercial polyclonal *S. aureus-specific* antibody from rabbits (specific antibody) and visualized bound antibodies by confocal laser scanning microscopy (CLSM, Figure 3A). As a control, we also labelled IgG_1_ specific to fluorescein isothiocyanate (anti-FITC IgG_1_) to evaluate unspecific binding by off-target antibodies to *S. aureus* cells, which could happen by interaction between the antibody’s Fc region and Protein A on the bacterial surface. Both planktonic cells and biofilms bound the nominally cognate antibody, while the off-target antibody was not detected (Figure 3A). The specific antibody was thus a suitable vehicle for delivering antimicrobials to *S. aureus* in suspension and within a biofilms.

**Figure 3.**
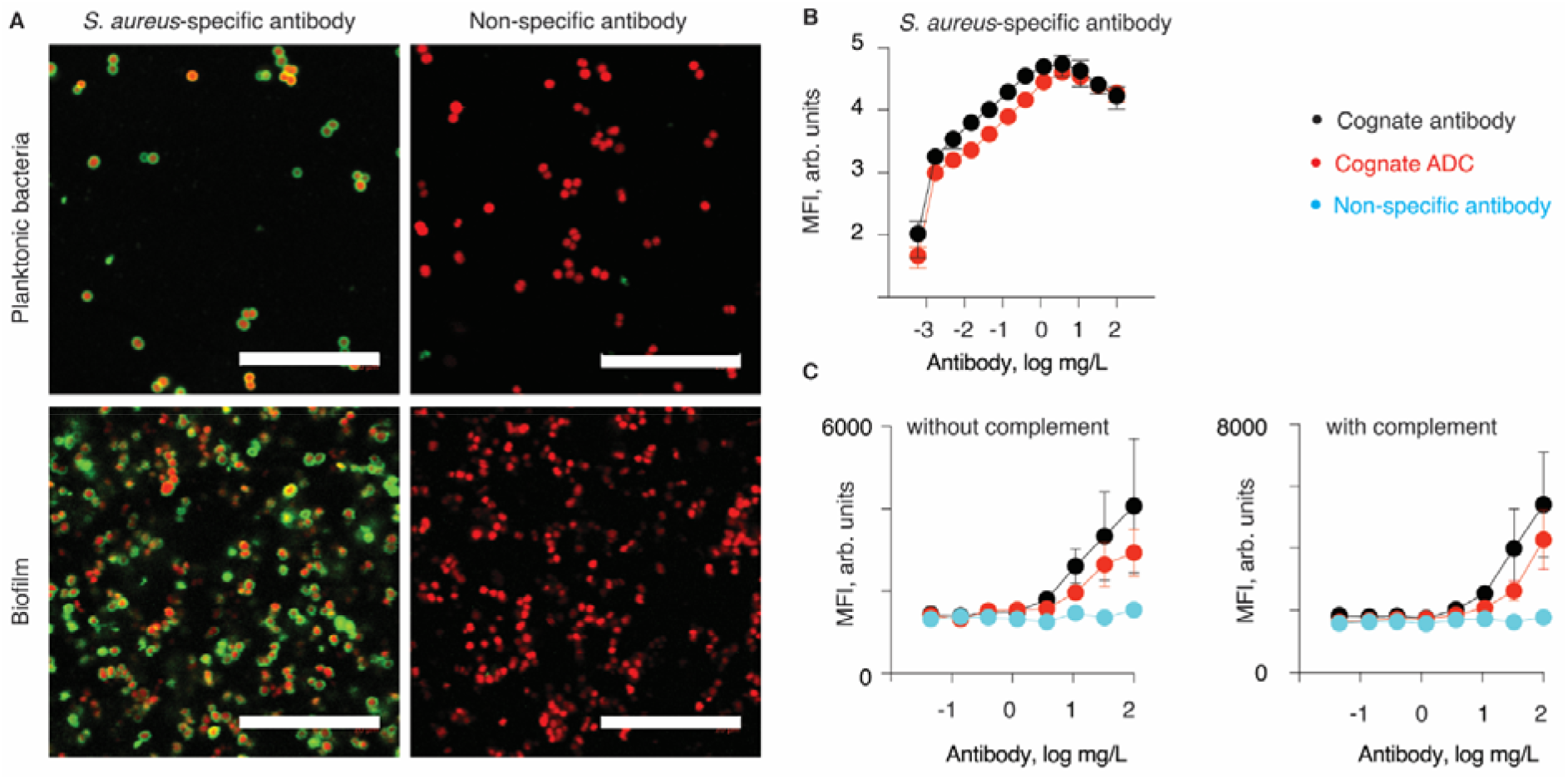
**(A)** CLSM images of *S. aureus* (red) following incubation with Alexa467-labelled *S. aureus-specific* antibody from rabbits or IgG_1_ specific to fluorescein isothiocyanate (anti-FITC IgG_1_) (green). Bacterial cells were visualized either by SYTO41 staining (biofilm) or GFP fluorescence (planktonic cells). Scale bars = 20 μm. **B)** Comparison of ADC (red) and the parent antibody (black) in their ability to bind to *S. aureus* strain Newman-spa/sbi-KO-mAm, quantified by flow cytometry as fluorescence from the bound secondary antibody.; **(c)** Specific/non-specific (black or blue, respectively) Antibody or ADC-mediated phagocytosis of *S. aureus* by human neutrophils over 15 min, quantified by flow cytometry measuring the GeoMFI of neutrophils with mAmetrine labelled *S. aureus* strain Newman Δspa/Δsbi_mAm. Each data point represents mean±SD of n=3 separate experiments.

Conversion of the antibody into the corresponding ADC may affect cognate interactions of the protein at both Fab (antigen recognition) and Fc (secondary interactions) regions. To test this, we first compared the *S. aureus-binding* properties of the parent antibody and the ADC constructed therefrom (Figure 3B). Antibodies or ADCs were incubated with *S. aureus* and thereafter fluorescently conjugated secondary antibody was added for the detection of bound protein. The parent antibody exhibited superior bacterial binding compared to the ADCs at the lowest tested concentrations (0.045-0.4 mg/L; p<0.009, two-way ANOVA). However, this was not the case at antibody concentrations used in the subsequent antimicrobial assays of this study (vide infra). Results of this experiment complement visual observations via CLSM (Figure 3A) and illustrate that the antibody and the ADC successfully bind the bacteria via cognate interactions.

From a different perspective, antibodies and ADC can also help to eliminate bacterial pathogens through opsonization, that is, binding to bacterium, possibly attracting complement proteins, and in doing so facilitating phagocytosis by neutrophils.^18,19^ This mode of action relies on the recognition between the Fc part of the antibody/ADC by the neutrophils and/or by the complement proteins, and this cognate interaction can possibly be altered by design of ADC. To investigate this, we used the Newman Δspa/Δsbi_mAm strain of *S. aureus* which expresses the fluorescent protein mAmetrine. Bacteria were incubated together with ADCs (specific and non-specific) or native antibodies, with or without 1% IgG/IgM depleted pooled human serum as complement source.^37^ Following this, the bacteria were incubated with human neutrophils, and the phagocytosis activity was quantified via flow cytometry (Figure 3C). In the absence of complement proteins, *S. aureus-specific* antibody and the ADC derived thereof promoted bacterial internalization by the neutrophils. In contrast, bacteria incubated with the non-specific antibody exhibited little, if any, internalization by the neutrophils. These data illustrate that both the cognate antibody and the cognate ADC bind to bacteria and facilitate internalization by neutrophils. The difference between the parent antibody and the ADC was statistically significant only at the highest protein concentration (10 mg/L) but not at the lower concentrations. Interestingly, addition of complement proteins afforded little change in the levels of bacteria internalization. This suggests that the cognate antibody and the ADC mediate phagocytosis via Fc gamma receptors. Together, results in Figure 3 illustrate that modification of the antibody with mitomycin C (at least up to a DAR of 8) had little effect on the cognate interactions via the Fab fragment with bacteria or via the Fc fragment with the neutrophils, which is highly important for the utility of the ADC as antimicrobial agents and potentially also as immune-stimulating agents.

### Antimicrobial efficacy of Mitomycin C

We chose mitomycin C as our antimicrobial agent due to its exceptional antimicrobial effect against biofilms, which otherwise have a high tolerance to antibiotics. To assess the concentration of mitomycin C needed to inhibit or kill *S. aureus*, we determined the minimum inhibitory concentration (MIC), the minimal biocidal concentration (MBC) required to kill > 99,9% of a planktonic culture, and the minimum biofilm eradication concentration (MBEC) required to fully eradicate viable cells from a biofilm (Table 1). The efficacy of antimicrobials differs slightly in various buffers and media, and we therefore determined the antimicrobial efficacy in all the solutions relevant for the subsequent antimicrobial testing. The similarity between MBC and MBEC values confirms that the antimicrobial efficacy of mitomycin C is not impacted by biofilm formation. Noteworthy, these values appear to be significantly lower than the reported values of the plasma level of mitomycin C that is reached in cancer therapy.^38^

**Table 1.** The antimicrobial efficacy of mitomycin C

| Media | MIC (mg/L) | MBC (mg/L) | MBEC (mg/L) |
| --- | --- | --- | --- |
| **BHI (complex media)** | 0.4 | 0.4 | 1-4 |
| **MM9 minimal media** | 0.4 | 0.4 | 1-2 |
| **MM9 buffer** | - | 0.1 | 1-8 |
| **MM9 buffer + NAC** | - | 0.1 | 1-4 |

### Antimicrobial efficacy of ADCs

The designed ADCs were tested as agents for delivery of antimicrobials to bacteria in suspension. To this end, *S. aureus* was incubated with the ADC for two h, and then diluted and plated on solid media to quantify viable cells as colony forming units (CFU). The ADC samples used in these experiments were designed using either specific pIgG (typical DAR = 7-8) or non-specific anti-FITC IgG_1_ (typical DAR = 13) and tested in a range of concentrations from 0.2 to 2 mg/L (equivalent mitomycin C concentration). Incubation of ADC with bacteria was performed in the presence or absence of NAC to trigger the release of mitomycin C. At the highest concentration tested (2 mg/L), incubation with both specific or non-specific ADC, with or without added NAC, decreased the number of viable bacteria to a value below the detection limit, likely indicating a non-specific drug release from all ADCs, even in the absence of NAC (Figure 4A). At lower concentrations, best seen at 0.5 mg/L, the antimicrobial activity of specific and non-specific ADCs was significantly different, and the cognate ADC decreased bacterial cell counts to below the detection level, whereas the nonspecific ADC only afforded a minor decrease. These data illustrate the highly desired outcome, namely that cognate ADCs exhibit pronounced therapeutic efficacy of antimicrobials at low concentrations by targeted delivery to the cell surface.

**Figure 4:**
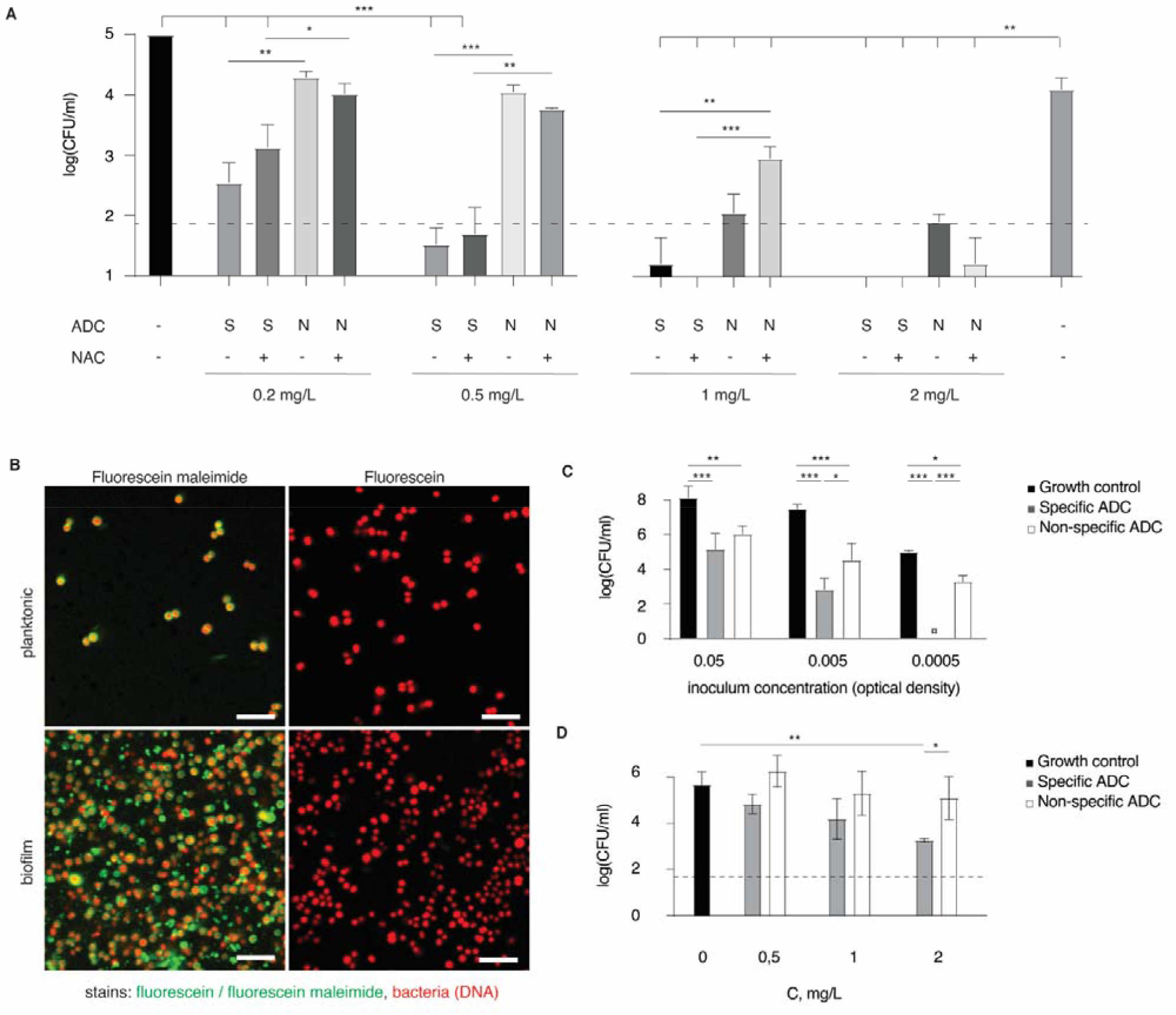
**a)** Quantification of live planktonic *S. aureus* in colony forming units after treatment with ADC and optionally with NAC; **b)** visualization of the surface accessible thiols on *S. aureus* (labelled with SYTO60, red) in planktonic state or within a biofilm using fluorescein maleimide or fluorescein (green) as a control. S scale bars= 10 μm; **c)** antimicrobial activity of the ADC (0.5 μg/ml) against planktonic bacteria, comparing the ADC cognate to *S. aureus* (“specific ADC”) or to FITC (“non-specific ADC”). Y axis shows viable *S.aureus* after 2 h incubation of ADC with different starting concentrations of bacteria. **d)** antimicrobial activity of the ADC against the *S.aureus* biofilms after 15 min incubation with ADCs to allow binding followed by 2 h incubation to allow ADCs to take effect before harvesting bacteria for CFU enumeration. Concentrations are expressed in equivalent concentration of mitomycin C. In panels a,c,d: Results are based on three biological replicates and presented as mean±st.dev.; statistical evaluation was conducted via a two-way ANOVA using log-transformed values of CFU count; for brevity, only selected statistical significance is presented; * *p* < 0.05; ** *p* < 0.01; *** *p* < 0.001.

The results in Figure 4A also demonstrate that NAC had no effect on the antibacterial activity of ADCs. This observation is highly un-expected. As such, it does not rule out that a NAC-mediated drug release occurred (as shown in Figure 1), but it indicates that the bacterial killing shown in Figure 4A is due to a drug release mechanism which is NAC-independent. One plausible mechanism can involve the thiol groups nested at the cell surface. For mammalian cells, exofacial sulfhydryls have become a highly important player in drug delivery at cell surfaces ^39^ and/or to the cell interior^28,29,34^. Thiols are also abundant on bacterial cells, ^31–33^ making it possible that interaction of ADCs with the cell surface ensues a thiol-disulfide exchange and release of the active drug. We therefore visualized thiol groups on the surface of *S. aureus* via fluorescent labelling. Bacteria were exposed to a thiol-reactive fluorescein maleimide or its non-reactive parent compound, fluorescein, and imaged by CLSM (Figure 4B). Exposure to fluorescein maleimide produced a strong fluorescent signal associated with the surface of bacteria, whereas no fluorescence was observed in the samples incubated with fluorescein. These data indicate that the *S. aureus* cell surface is rich in accessible, reactive thiol groups.

In designing the next experiment, we considered that the interaction between bacteria and ADC, both specific and non-specific, is concentration-dependant, and that antimicrobial efficacy of ADC should therefore be dependent on the concentration of bacteria in suspension. Indeed, in our hands, the antimicrobial activity of the ADC was strongly dependent on the bacterial cell concentration (Figure 4c). Moreover, the bacterial cell concentration was highly important in defining the relative efficacy of antimicrobial activity between the cognate ADC and its non-specific counterpart. At the highest cell concentration, bacterial cell killing was strong for both specific and non-specific ADC preparations, and the difference between the two ADCs was not significant. In contrast, at the lowest bacterial cell concentration, the efficacy of treatment with the cognate ADC was significantly higher than that for the non-specific ADC. This experimental finding is readily explained by the difference in potency between the two ADCs. The low-potency non-specific interactions may be significant at a high cell concentration, but they should become less significant at the low cell concentrations. At the same time, the high potency cognate interactions remain pronounced within the studied range of cell concentrations, and this leads to an increasing difference between specific and non-specific ADC by efficacy of antimicrobial activity.

Having established the antimicrobial effect of ADCs against planktonic *S. aureus*, we also aimed to validate if this mode of drug delivery is effective against the bacterial biofilms (Figure 4d). The antimicrobial effect on biofilms was concentration-dependent, and at 2 mg/L concentration of ADC, the antimicrobial effect of specific ADCs was significantly higher than the non-specific ADCs. The biofilm was not eliminated during the treatment, but it should be stressed that the treatment time was very short. Biofilms were only exposed to ADCs in solution for 15 min and then incubated for 2 h after unbound ADC was removed. Even after this short incubation time, the number of viable bacteria decreased by more than 100-fold, illustrating that ADCs are efficacious against *S. aureus* biofilms.

### Therapeutic efficacy of ADCs against biofilm infections

*In vivo* evaluation of ADC was performed in 8-10 weeks old C57bl/6j mice, in an implant-associated osteomyelitis model (Figure 5a). ^40^ Briefly, stainless steel insect pins were surgically inserted into the tibia after they had been inoculated in an overnight culture of *S. aureus* SAU060112 (Ref. ^41^). Control mice received a sterile implant. On day 3 after surgery, mice were administered with fluorescently labelled antibodies to verify the ability of the cognate antibody to accumulate at the site of infection. For comparison, fluorescently labelled anti-FITC antibodies were administered to evaluate non-specific accumulation of antibodies. *In vivo* full body imaging was used to visualize and quantify fluorescence from antibodies in the mice with sterile compared to infected implants, and from the specific antibodies compared to non-specific antibodies (Figure 5B,C). 24 h after antibody administration, fluorescence from *S. aureus-specific* antibodies was higher from infected implants compared to sterile implants (marked in Figure 5C with “+” and “-” respectively), whereas the signal for the non-specific antibody was the same from the infected and the sterile implants. These data indicate that the *S. aureus-specific* antibody bound to and accumulated at the site of infection, as is required for the desired site-specific therapeutic activity.

**Figure 5.**
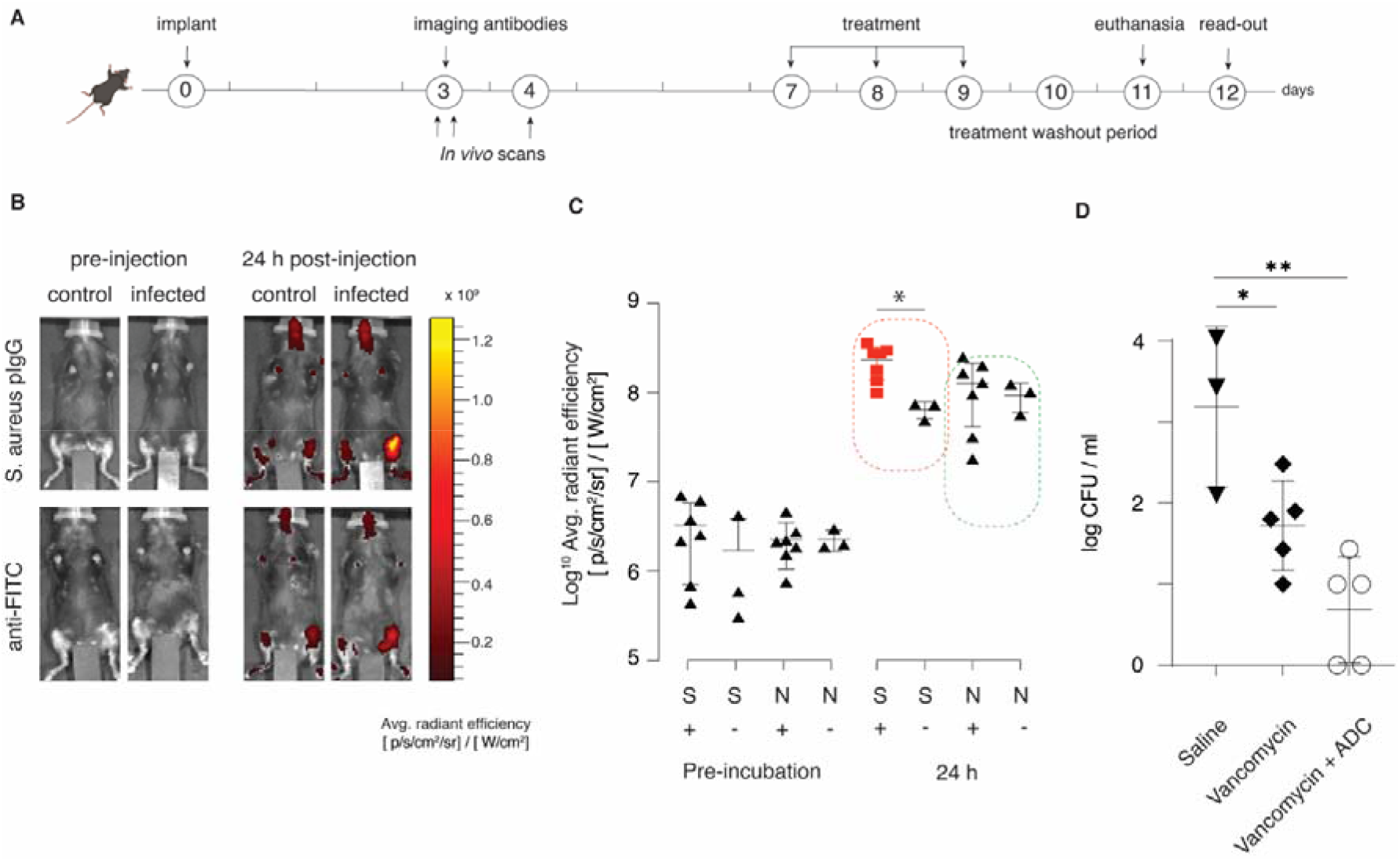
Pilot imaging and therapy in vivo study. **(a)** schematic presentation of the experiment timeline; **(b)** *in vivo* imaging of deposition of the fluorescently labelled antibodies, cognate to *S. aureus* or to fluorescein, in a limb with a sterile implant or in the limb with an infected implant (osteomyelitis model); **(c)** quantification of deposition of the fluorescently labelled antibodies; S = *S. aureus* specific antibody; N = non-specific antibody; in both cases labelled with Alexa Fluor NHS ester 687; “+” signifies animals with an infected implant, “-“ is for animals with a sterile implant; **(d)** antimicrobial effects in mice upon treatment with vancomycin, ADC (cognate to *S. aureus)*, or the combination of two agents. In panels c,d:: each data point represents radiant efficiency from implanted leg (C) or represents a single animal (D); statistical evaluation was conducted via two-way ANOVA (panel **c**) or one-way ANOVA (panel **d**).

Seven days after the surgery, animals with infected implants were randomly divided into three treatment groups, that were administered with saline, vancomycin as a monotherapy, or vancomycin combination therapy with ADC, for three consecutive days. This pilot experiment was designed specifically, such that the two treatment arms contained vancomycin, which is first line treatment of MRSA infection. ^42^ The administered dose of vancomycin was 110 mg/kg/12h/s.c.. For the ADC (DAR 7-8), the administered dose was 5 mg / kg / 24h / i.v.. Here and in all *in vivo* experiments, ADC concentration is expressed in total solids content (not equivalent concentration of mitomycin C); 5 mg/kg ADC corresponds to approx. 4.9 mg/kg of the antibody and 86 μg/kg of mitomycin C. Treatments were administered on days 7, 8, and 9 and followed by a two-day washout period before animals were euthanized, and the bacterial load was quantified by CFU enumeration. Quantification of bacteria on the surface of the recovered implant revealed that vancomycin alone led to an appr. 10-fold decrease in the bacterial load (Figure 5D) while addition of ADC to the vancomycin treatment led to an appr. 100-fold reduction, although with a limited sample size the effect was not statistically significant. Nevertheless, this pilot study was important in that it revealed that the ADC treatment did not have any detrimental effect to animal well-being, and it also provided the first indication of therapeutic efficacy of the ADC *in vivo*.

Next, we performed two experiments to investigate the therapeutic potential of the ADC. First, ADC mono-therapy was compared to mono-therapy with vancomycin or a combination-therapy with vancomycin and ADC (Figure 6A,B). In this experiment, unlike the pilot study discussed above in Figure 5, vancomycin treatment alone afforded no effect on the bacterial burden, which is explained by the differences in the implant inoculation and the implant recovery protocols employed in the two experiments. In stark contrast, ADC treatment was highly effective and statistically significant compared to saline treatment and the vancomycin treatment arms. Concurrent administration of vancomycin and the ADC afforded no added benefit for the therapeutic outcome. These data provide a strong indication of efficacy of treatment with the developed ADC *in vivo*, even after a very short (relative to the clinically standard, Ref. ^43,44^) treatment period of three single doses of the ADC.

**Figure 6.**
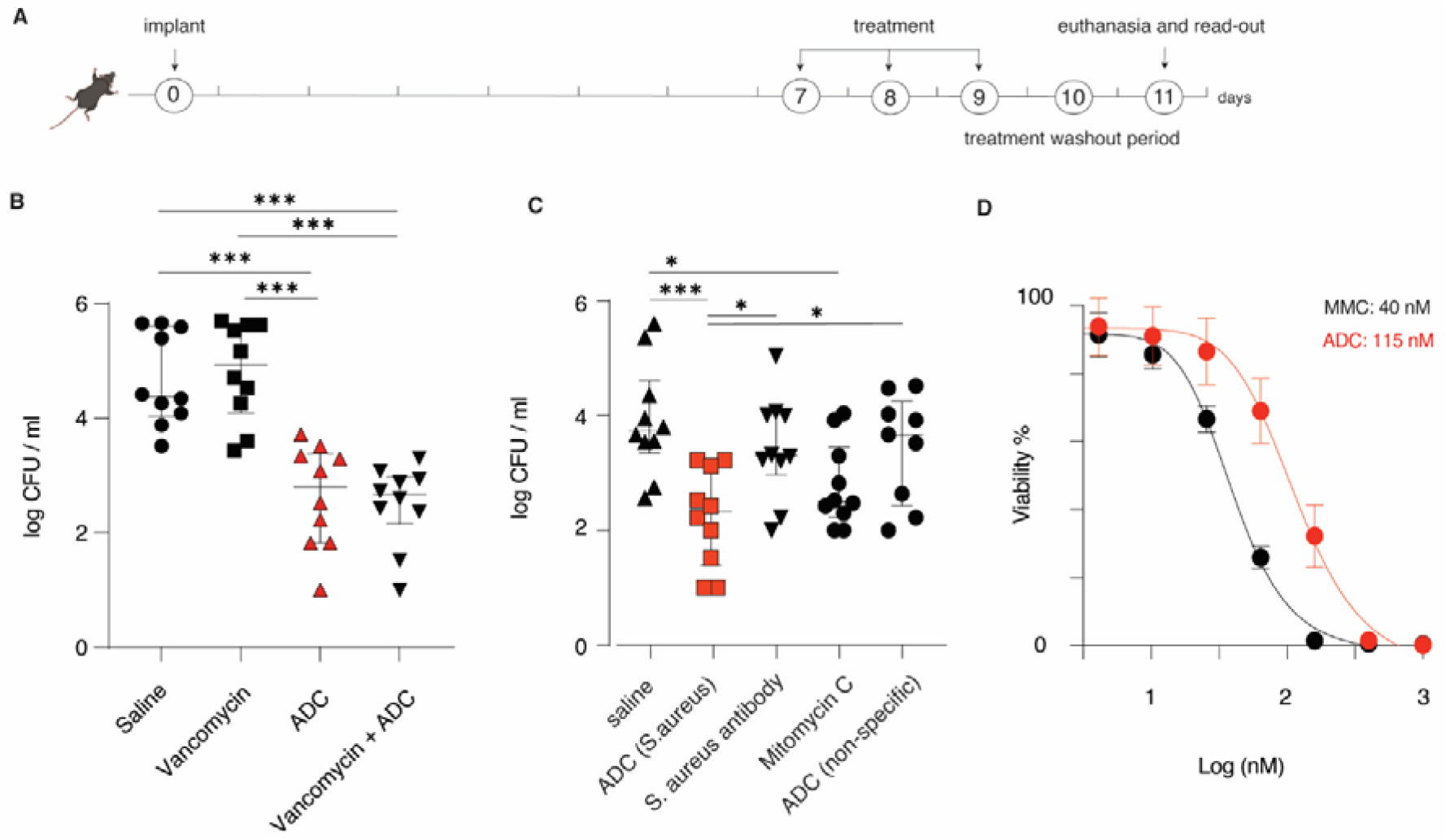
**(A)** Schematic representation of the work flow timeline for the quantification of antibacterial effects in vivo; **(B)** experimental antimicrobial effects upon treatment with vancomycin (110 mg/kg/12h/s.c.), ADC (5 mg/kg/24h/i.v.), or the combination of the two agents **(C)** experimental antimicrobial effects upon treatment with ADC (cognate to *S. aureus* or fluorescein, in both cases 5 mg/kg/24h/i.v.), with the anti-*S. aureus* antibody (5 mg/kg/24h) or mitomycin (86 μg/kg/24h/iv) taken individually; in panels **B,C**: each data point represents an individual mouse; statistical significance was calculated via a one-way ANOVA using the log-transformed experimental CFU values. **(D)** In vitro cytotoxicity of mitomycin and an ADC derived thereof for MOLT-4 cells (a human T lymphoblast cell line) following a 72 h incubation (presented results are based on three independent experiments and presented as mean ± std.dev.) *P≤0.05 ***P≤0.001

Lastly, efficacy of the treatment was quantified side by side for the specific and non-specific ADC designs based on antibodies cognate to *S. aureus* or off-target antibodies, and also to the cognate antibody or mitomycin C taken individually (Figure 6c). The most important observation from this experiment is that the therapeutic benefit of the specific ADC cognate to *S. aureus* was statistically significant compared to the treatment with saline, to the treatment with the unconjugated antibodies, and to the treatment with non-specific ADC. The superior therapeutic effect of the *S. aureus*-specific ADC supports our presumption that the bacteria are killed by mitomycin C released from the ADCs upon interaction with the biofilm. Mitomycin C monotherapy had also a statistically significant effect, which is consistent with the antibacterial efficacy that we and others have observed for the drug in bacterial cell culture. ^35,45^ Comparison between the specific ADC and mitomycin C monotherapy revealed no statistical significance. Nevertheless, *in vitro* cell culture experiments illustrate that ADC is less toxic when compared to mitomycin C (Figure 6D), which implies that the ADC treatment may be associated with fewer side effects.

## Discussion

In this study, we developed an ADC that targets *S. aureus* and exhibits antimicrobial activity *in vitro* and *in vivo*. To our knowledge, this is the first study to develop and demonstrate the potential for using ADCs in antimicrobial therapy directed at extracellular infections and biofilms. Only few prior studies have developed ADCs for antimicrobial therapies, ^20^ and the focus was mainly been on intracellular drug release, to kill bacteria internalized by immune cells.^19,46^ Effective antimicrobial therapies against biofilm infections typically require a high load of antibiotics and lengthy treatment.^47,48^ Enhanced deliverable payload can be achieved using tools of nanomedicine, such as liposomes or solid nanoparticles, ^10,49^ whereas enzymes and nanozymes may prove useful for localized drug syntheses schemes^12,50^. Each of these techniques has its own merits. ADC have a competitive edge in that multiple products are already on the market and numerous candidates navigate through clinical trials, illustrating that technology for the production of ADC is well-established, and clinical acceptance of these agents is very high.^14,15^ However, as described to date, ADC focus on cancer treatment and intracellular drug release.^14,15^ For the treatment of bacterial pathogens and specifically biofilms, a novel mode of action for ADC is required. Our initial thought was to use ADC for drug targeting and thereafter achieve a localized drug release using an independently administered trigger for drug release. Somewhat surprisingly, we observed that this trigger was not required and ADC proved to be potent and efficacious as a monotherapy, *in vitro* and *in vivo*.

The ADC synthesized in this work release the payload via disulfide reshuffling. We validated via thiol staining that the bacterial cell surface has abundant thiols, which strongly suggests that bacteria are competent to participate in the thiol-disulfide exchange, to initiate the drug release from the ADC. Indeed, disulfide reshuffling has been documented at the surface mammalian cells in numerous studies.^28–30,34^ For bacteria, such reports are scant and in fact antibiotics acting in the cytoplasm were inactivated via their conversion to disulfide-containing derivatives, possibly suggesting conjugation to the bacterial cell surface and thereby arrested drug cell entry.^31^ In recent studies, Shchelik and Gademann^51,52^ observed that the disulfide containing derivatives of vancomycin and cephalosporin were superior to the parent antibiotic molecules, although no mechanism for the involvement of the disulfide functionality was provided to explain the enhanced drug efficacy of the newly synthesized compounds. In our hands, attempts to block the bacterial cell surface thiols with maleimide-based sulfhydryl poisons ^30^ were met with limited success (data not shown). Thus, detailed understanding of the mechanism of activity of the ADC and the role of bacteria-mediated drug release requires significant further experimentation. Nevertheless, all the data collected in this work point towards the bacteria-mediated drug release because: i) spontaneous drug release from the ADC in the MM9 culture media was insignificant; ii) external triggering of drug release by added NAC did not enhance antimicrobial activity; and iii) cognate ADC exhibits superior efficacy and potency compared to the non-specific counterpart.

We believe that the results of this study are highly important in that these opens doors for using mitomycin C as an antimicrobial agent. Mitomycin C works by crosslinking DNA and is thus highly cytotoxic. It is a highly attractive agent for the use as an antimicrobial because it is equally effective against actively growing cells (susceptible to conventional antibiotics) and slow-growing or dormant persister cells (tolerant to conventional antibiotics). A number of studies have confirmed the potential of mitomycin C as a potent antimicrobial against biofilms due to its effect on persister cells ^35,45,53^. Our study confirms that this potency translates to a superior treatment outcome for biofilm infections, specifically, when administered as a targeted formulation for localized drug release within the biofilm.

## Conclusions

Antibody-drug conjugates, one of the most successful platforms for targeted intracellular drug delivery, are developed in this study towards a new area of applications, specifically targeted delivery of drugs to treat bacterial biofilms. Our results point to a novel mechanism for drug release at the bacterial cell surface, namely via a thiol-disulfide reshuffling between the disulfide-conjugated drug and the exofacial cellular thiols. We demonstrate antimicrobial efficacy of the ADC *in vitro* and *in vivo* in an implant-associated osteomyelitis model. Treatment of biofilms is a tremendous socio-economic burden and represents an unmet medical need. We believe that our findings open up significant opportunities to the treatment of bacterial biofilms, with a potential to identify ligands for optimized targeting and/or therapeutic molecules for enhanced antimicrobial effects.

## Acknowledgements

This work received financial support from the Independent Research Fund Denmark (Project No. FTP-9041-00242A, to ANZ), and the Novo Nordisk Foundation (Project No. NNF19OC0058357). This publication is part of the project DARTBAC (project number NWA.1292.19.354 of the research programme NWA-ORC which is partly financed by the Dutch Research Council, NWO).

## TOC image

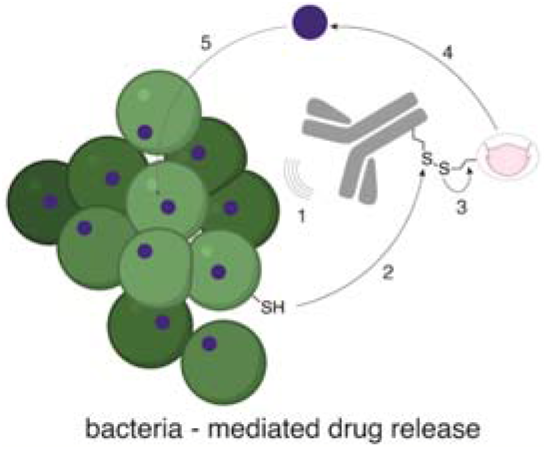

## Synopsis

New mode of action for the antimicrobial agents are urgently required. We present bacteria-mediated drug release from antibody-drug conjugates, active against biofilms in vivo.

