## Supplementary information for "Antibody-drug conjugates to treat bacterial biofilms"

**Materials and methods:**

**General information:**

All chemicals and reagents were purchased from commercial vendors (Sigma Aldrich, Tokyo Chemical Industry, Selleck chemicals) and used without further purification, unless otherwise stated. Triethylamine (TEA) and *N,N*-dimethylformamide (DMF) was purchased anhydrous and the deuterated solvents were purchased from EurisoTop. Dry dichloromethane (DCM) were collected from a MBraun SP800 purification system. The remaining solvents were HPLC graded. Moisture sensitive reaction were performed in flame-dried glassware under positive pressure of N_2_. Analytical thin layer chromatography (TLC) was performed using pre-coated aluminum-packed plates (Merk® Silica gel 60 F_254_) and visualized under UV irradiation (254 nm or 365 nm) and/or dipping into KMnO_4_- or Ninhydrin stain followed by gently heating. Flash column chromatography was carried out using silica gel – high purity grade (w/Ca, ~0.1%, 230-400 mesh particle size, 60Å pore size) as the stationary phase acquired by Sigma-Aldrich. Ultraviolet-visible (UV-vis) absorbance spectra were measured using a Thermo Scientific Nanodrop 2000c.

Nuclear Magnetic Resonance (NMR) spectra were recorded on a Varian Mercury 400 MHz spectrometer and the spectra were recorded either as ^1^H-NMR 400 MHz or ^13^C-NMR 101 MHz. All spectra were referenced to the solvent peak; the chemical shifts are reported in ppm relative to the residual solvent peak. Analysis of the spectra was performed using MestReNova. The following abbreviations are used to indicate the multiplicity in the NMR spectra: s = singlet, d = doublet, t = triplet, q = quartet, dd = double doublet, m = multiplet. Coupling constants are reported in hertz (Hz) as the mean value between coupled hydrogen atoms. ^13^C-NMR spectra were acquired in a broadband decoupled mode. High-resolution mass spectrometry (HR-MS) was recorded on a Bruker Maxis Impact TOF-MS using electrospray ionization (ESI+). The spectra were calibrated to an internal standard and analyzed with the DataAnalysis software.

Matrix assisted laser desorption ionization-time-of-flight mass spectroscopy (MALDI-TOF MS) measurements were recorded using a Bruker AutoFlex II MS with nitrogen laser (337 nm) and 20 kV accelerating voltage with a grid voltage of 90%. At least 100 laser shots covering the complete spot were accumulated for each spectrum. For drug-to-antibody ratio of the prepared conjugates, sinapinic acid (20 mg/mL) in 50% acetonitrile with 0.1% trifluoroacetic acid was used as matrix. Antibody solution (1.0 mg/mL) was mixed with an equal volume of matrix, and 2 μL of the resulting mixture was loaded onto MTP 384 ground steel target plate. The extent of modification was determined by subtracting the antibody-drug conjugates’ m/z values from the native antibody m/z and dividing by the molecular weight of mitomycin linker (542 g/mol).

Analytical High-Performance Liquid Chromatography (HPLC) analysis was carried out an Agilent 1260 Infinity II with an Agilent ZORBAX eclipse Plus C18 column with a particle size at 3.5 𝜇m, a length at 150 mm and an internal diameter of 4.6 mm. Mobile phases were ultrapure water (MQ) and acetonitrile that was cooled overnight in the fridge to 4 ^o^C and kept on ice during the HPLC analysis. MQ was received from Milli Q direct 8 system (Millipore). Furthermore, all the samples were kept on ice (0 ^o^C) until analysis to avoid side-degradation reactions.

**Syntheses.**

**Compound 1b.** 2-hydoxyethyldisulfide (300 mg, 1.94 mmol, 1 equiv.) was dissolved in CH_2_Cl_2_ (16 ml) and subsequently addition of 4-nitrophenyl chloroformate (871 mg, 4.32 mmol, 2.23 equiv.). The mixture was bubbled through with N_2_ and cooled to 0 ^o^C followed by dropwise addition of TEA (1.08 ml, 7.76 mmol, 4 equiv.). After addition the reaction was left to heat to room temperature and stirred for two h. The reaction was quenched with NH_4_Cl, washed with brine, dried over Na_2_SO_4_ and concentrated *in vacuo*. The crude was purified with flash column chromatography (1:1 pentane:CH_2_Cl_2_ to 3:7 pentane: CH_2_Cl_2_) yielding the pure product as a yellow powder (467 mg, 0,964 mmol, 50 % and 61 %). **^1^H NMR** (400 MHz, Chloroform-d) δ 8.28 (d, J = 9.2 Hz, 4H), 7.39 (d, J = 9.2 Hz, 4H), 4.57 (t, J = 6.5 Hz, 4H), 3.08 (t, J = 6.5 Hz, 4H). **^13^C NMR** (101 MHz, Chloroform-*d*) δ 155.47, 152.48, 145.66, 125.50, 121.91, 66.89, 36.90. **HRMS** (ESI+) *m/z* calculated (calcd.) for C_18_H_16_N_2_O_10_S_2_ + Na^+^: 507.0139; found: 507.0173, calcd. for C_18_H_16_N_2_O_10_S_2_ + K^+^: 522.9978; found: 522.9924, calcd. for 2C_18_H_16_N_2_O_10_S_2_ + Na^+^:991.0384; found: 991.047

**Compound 1. 1b** (22.2 mg, 0.0458 mmol, 2.4 equiv.) was dissolved in dry DMF (0.5 mL) and stirred under N_2_. In another flask mitomycin C (6.4 mg, 0.01914 mmol, 1 equiv.) and TEA (3.99 µL, 0.0287 mmol, 1.5 equiv.) was dissolved in dry DMF (0.25 mL) followed by dropwise addition to the stirring reaction mixture. HOBt (17.2 mg, 0.127, 6.7 equiv.) was subsequently added and the reaction was left in dark for 24 h, concentrated and purified by flash column chromatography (1:1 pentane:EtOAc to 3:7 pentane:EtOAc) to yield the product as a purple powder (7.4 mg, 0.011 mmol, 57 %). R_f_ (3:7 Pentane:EtOAc) = 0.26. **^1^H NMR** (400 MHz, Chloroform-*d*) δ 8.29 (d, *J* = 9.2 Hz, 2H), 7.40 (d, *J* = 9.2 Hz, 2H), 4.84 (dd, *J* = 10.8, 4.7 Hz, 1H), 4.53 (t, *J* = 6.6 Hz, 2H), 4.39 – 4.26 (m, 3H), 3.68 (dd, *J* = 11.1, 4.7 Hz, 1H), 3.49 (dd, *J* = 13.4, 1.9 Hz, 1H), 3.44 (d, *J* = 4.6 Hz, 1H), 3.33 (dd, *J* = 4.6, 1.8 Hz, 1H), 3.20 (s, 3H), 3.03 (t, *J* = 6.6 Hz, 2H), 2.96 (t, *J* = 6.7 Hz, 2H), 2.89 (t, *J* = 5.7 Hz, 1H), 1.76 (s, 3H). **^13^C NMR** (101 MHz, CDCl_3_) δ 160.74, 156.42, 155.51, 154.43, 152.51, 147.09, 125.52, 121.99, 110.57, 105.58, 105.41, 66.98, 64.74, 62.43, 60.51, 49.94, 48.79, 43.70, 41.97, 41.37, 40.13, 36.80, 36.76, 14.37, 8.05. **HRMS** (ESI+) *m/z* calculated (calcd.) for C_27_H_29_N_5_O_12_S_2_ + H^+^: 680.1328, found: 680,1328; C_27_H_29_N_5_O_12_S_2_ + Na^+^: 702.1146, found: 702.1149; C_27_H_29_N_5_O_12_S_2_ + K^+^: 718.0886, found: 718.0891.

**Compound 2b.** 3-mercapto-3-methylbutan-1-ol (200 mg, 1.66 mmol, 1 equiv.) was dissolved in CH_2_Cl_2_:EtOAc (1:1, 52 mL in total). CuCl_2_ (1.120 g, 8.33 mmol, 5 equiv.) was dissolved in 0.5 M HCl (36 mL) followed by addition to the organic phase. The reaction was left stirring for 1.5 h at r.t. followed by extraction of the reaction mixture with CH_2_Cl_2_ and afterwards of the aqueous phase. The organic phases were combined, washed with brine, dried with Na_2_SO_4_, filtered and concentrated yielding the pure product as a clear oil (289.3 mg, 1.212 mmol, 73 %). **^1^H NMR** (400 MHz, Chloroform-*d*) δ 3.81 (t, *J* = 7.1 Hz, 4H), 1.86 (t, *J* = 7.1 Hz, 5H), 1.32 (s, 13H). **^13^C NMR** (101 MHz, Chloroform-*d*) δ 59.85, 48.55, 44.14, 28.82. **HRMS** (ESI+) *m/z* calculated (calcd.) for C_10_H_22_O_2_S_2_ + Na^+^: 261.0953, found: 261.0950; Cald. for C_10_H_22_O_2_S_2_ + 2Na^+^-H^+^: 283.0772, found: 283.0708

**Compound 2c. 2b** (86.7 mg, 0.364 mmol, 1 equiv.) was dissolved in dry CH_2_Cl_2_ (0.75 ml) followed by addition of TEA (125 µL, 0.895 mmol, 2.5 equiv.). In another flask 4-nitrophenyl chloroformate (207 mg, 1.03 mmol, 2.82 equiv.) was dissolved in dry CH_2_Cl_2_ (0.75 ml), cooled to 0 ^o^C and the mixture containing TEA and 2-hydroxytheyldisulfide was added dropwise to the stirring mixture. After addition, the ice was removed, and the reaction mixture was left stirring as a slurry at room temperature for three h.  The reaction was diluted with CH_2_Cl_2_ (8 ml) and quenched with NH_4_Cl, washed 5 times with brine and dried over Na_2_SO_4_. The crude was concentrated yielding the product as a pale yellow oil (142.1 mg, 0.25 mmol, 69 %). **^1^H NMR** (400 MHz, Chloroform-*d*) δ 8.28 (d, *J* = 9.1 Hz, 4H), 7.38 (d, *J* = 9.1 Hz, 4H), 4.43 (t, *J* = 7.1 Hz, 4H), 2.05 (t, *J* = 7.1 Hz, 4H), 1.37 (s, 12H). **^13^C NMR** (101 MHz, Chloroform-*d*) δ 155.62, 152.55, 145.54, 125.46, 121.94, 66.67, 48.12, 40.24, 28.64. **HRMS** (ESI+) *m/z* calculated (calcd.) for C_24_H_28_N_2_O_10_S_2_ + Na^+^: 591.1077, found: 591.1083; Cald. for C_24_H_28_N_2_O_10_S_2_ + K^+^: 607.0817, found: 607.0885;

**Compound 2. 2c** (38.6 mg, 0.0678 mmol, 2 equiv.) was dissolved in dry DMF (1 mL) and stirred under N_2_. In another flask mitomycin C ( 11.35 mg, 0.0339 mmol, 1 equiv.) and TEA (7.09 µl, 0.0508 mmol, 1.5 equiv.) was dissolved in dry DMF (0.5 mL) followed by dropwise addition to the stirring reaction mixture. HOBt (27.5 mg, 0.203 mmol, 6 equiv.) was subsequently added and the reaction was left in dark for 24 h, concentrated and purified by flash column chromatography (1:1 pentane:EtOAc to 3:7 pentane:EtOAc) to yield the product as a purple powder (13.43 mg, 0.0176 mmol, 52 %). R_f_ (3:7 Pentane:EtOAc) = 0.31. **^1^H NMR** (400 MHz, Chloroform-*d*) δ 8.28 (d, *J* = 9.1 Hz, 2H), 7.38 (d, *J* = 9.2 Hz, 2H), 4.88 (dd, *J* = 10.8, 4.7 Hz, 1H), 4.48 – 4.36 (m, 3H), 4.31 – 4.15 (m, 3H), 3.67 (dd, *J* = 11.0, 4.7 Hz, 1H), 3.49 (dd, *J* = 13.4, 1.9 Hz, 1H), 3.42 (d, *J* = 4.6 Hz, 1H), 3.29 (dd, *J* = 4.6, 1.8 Hz, 1H), 3.19 (s, 3H), 2.01 (t, *J* = 7.1 Hz, 2H), 1.90 (t, *J* = 7.3 Hz, 2H), 1.76 (s, 3H), 1.34 (s, 6H), 1.29 (s, 6H). **^13^C NMR** (101 MHz, CDCl_3_) δ 178.58, 176.05, 160.95, 156.36, 155.68, 154.42, 152.55, 147.15, 145.58, 125.46, 121.96, 110.71, 105.63, 105.35, 77.16, 66.73, 64.35, 62.31, 49.90, 48.85, 48.25, 48.01, 43.58, 42.12, 40.25, 40.14, 28.63, 28.62, 28.58, 8.03. **HRMS** (ESI+) *m/z* calculated (calcd.) for C_33_H_41_N_5_O_12_S_2_ + H^+^: 764.2266, found: 764.2288; Cald. for C_33_H_41_N_5_O_12_S_2_ + Na^+^: 786.2085, found: 786.2133;

**Fluorescence labelling antibodies.** Anti *S. aureus* antibody (Catalog nr.:PA1-7246) and anti-FITC (Catalog nr.: 31242) and Alexa Flour^TM^ NHS ester 680 antibody were purchased from ThermoFisher. Protein concentrations were determined using UV-vis (λ_max_= abs_280 nm_) with ε_protein_= 210,000 M^-1^cm^-1^. Solutions of antibodies were buffer exchanged by the use of spin filters (Amicon spin filters, 30 kDa cutoff, regenerated cellulose) to 0.1 M sodium bicarbonate buffer, pH 8.3 and adjusted to a final protein concentration of >3 g/L . The fluorophore (Alexa Flour 680 NHS ester or Alexa Flour 647 NHS ester) was dissolved in DMSO (10 g/L final concentration) immediately before use and added to an antibody solution in an excess of 8 molar equivalents. The reactions were left at room temperature for one hour under mild shaking (800 rpm) in the dark. The solutions were then buffer exchanged to 10 mM PBS, pH 7.4 by the use of spin filters (Amicon spin filters, 30 kDa cutoff, regenerated cellulose) followed by purification by gel filtration through a NAP-5 column (Sephadex G-25 DNA grade).

**Synthesis of ADC.** General protocol: solutions of an antibody was buffer exchanged into a 10 mM PBS buffer, pH 7.4, using Amicon spin filters (MW cutoff 30 kDa, regenerated cellulose), with a final protein concentration at > 3 g/L. Compound **1** (55 equiv. to protein) was added and the reaction was left shaking (800 rpm) in the dark at room temperature for 2 h. The ADC was purified with gel filtration through a NAP-5 column (Sephadex, G-25 DNA grade). The DAR was determined based on the distinct absorbance for mitomycin C at 360 nm and the absorbance at 280 nm for the protein.

**Drug release from ADC.** Triggered drug release was tested using solutions of GSH, DTT, NAC (each taken at 5 mM concentration), and BSA (0.76 µM) in PBS; unspecific drug release/linker stability was tested in PBS and in two different types of cell media, namely RPM1-1640 medium containing 10 % FBS, 1% P/S, 2 mM L-Gln (denoted as FBS) and Modified M9 buffer (mM9). All the samples were incubated at 37 $^{\circ}$C and at each time point (15 minutes, 1 hour, 24 h,) an aliquot was draw out and analysed on HPLC.

***In vitro* bacterial work:**

**Bacterial strains and growth conditions.** Bacterial cultures were stored in 15 % glycerol at -80 °C. Single colonies were grown on brain heart infusion (BHI) agar and stored at 4-8 °C. Experiments were performed with overnight cultures. Each biological replicate was inoculated from a single colony in BHI broth at 37 °C, 180 rpm. For antibody binding and phagocytosis activity measured by flow cytometry we used *S. aureus* Newman Δspa/Δsbi_mAm, which as low or no expression of staphylococcal protein A (SpA) and second immunoglobulin-binding protein (Sbi)^54^, and contains a plasmid from which the fluorescent protein mAmetrine (mAm) is constitutively expressed. ^55^ For antibody binding measured by microscopy, we used *S. aureus* ATCC29213 transformed with the plasmid pSB2019,^56^ which expresses green fluorescent protein (GFP) constitutively. Plasmid maintenance was secured by growing and incubating cells in BHI media containing 10 μg/mL chloramphenicol. *S. aureus* ATCC 29213 was used for all other *in vitro* experiments. All experiments were performed with at least 3 biological replicates of independently grown overnight cultures.

**Media and buffers.** The growth media used in the experiments was Brain Heart Infusion Broth (BHI). Modified M9 buffer (mM9) contained M9 minimal salts 5 X (Sigma-Aldrich M6030), 2 mM MgSO4, 0.1 mM CaCl2, 1 mM Thiamine-HCL, 0.05 mM nicotinamide and 1 mL/L trace metals (TMS3). The pH of mM9 was adjusted to 7.4. After autoclavation components were sterile filtered and added to the buffer. mM9 media with carbon and casamino acids (mM9 + carbon) was supplemented with 1 % glucose and 1 % casamino acids. Assays involving mM9 with N-acetyl cysteine (NAC) contained 5 mM NAC.

**MIC, MBC and MBEC of mitomycin C.** A two-fold serial dilution of mitomycin C (from 0.00625 - 3.2 mg/L) was prepared in 96-well plates in mM9 buffer, mM9 buffer with 5 mM NAC, mM9 media (mM9 buffer with 1 % glucose and 1 % casamino acids), or BHI. Overnight cultures were harvested by centrifugation (13150 x g rpm, 8 min) and resuspended to OD_600_ of 0.05 in media matching the microtiter pates and inoculated into the plates by 10 fold dilution, and incubated at 37 °C for 19 h 180 rpm before measuring OD_600_ (BioTek,Powerwave XS2). MIC was defined as the lowest concentration resulting in >90% inhibition of growth. MBC was determined by spotting out 10 μL from wells with no apparent growth on BHI agar and incubating for 24 h at 37 °C. MBC was determined as the lowest concentration resulting in no detection of viable cells (corresponding to a >99,9% reduction in CFU).

To determine MBEC, biofilms were grown on peg-lids (NuncTM 445497 ImmunoTM TSP Lids) by inoculating the pegs in an overnight culture for 30 min at 37 °C and transferring to BHI for biofilm growth at 37 °C for 24 h with 120 rpm shaking. MBEC was determined in mM9 buffer with and without 5 mM NAC, mM9 media, and BHI. A two-fold serial dilution of mitomycin C was prepared for these media (concentration range 128 - 0.25 mg/L), and peg lids with biofilms incubated in these plates for 24 h at 37 °C and 120 rpm. After treatment, biofilms were washed by dipping peg lids twice in 96-well plates with BHI to remove mitomycin C, and peg lids were then placed in a recovery plate with BHI, sonicated for 10 min (USCS1700 T, VWR, Westchester, PA, USA) to remove the biofilm from the peg-lids, and incubated for 72 h at 37 °C, 50 rpm. Growth in the recovery plate was detected by measuring OD_600_ and the MBEC was determined as the lowest concentration resulting in no growth.

**Antibody binding to *S. aureus* detected by microscopy*.*** Planktonic overnight cultures of *S. aureus* ATCC29213_GFP were immobilised by adsorption to SuperFrost Ultra Plus Adhesion Slides, blocked with 1% BSA, rinsed 3x with mM9 buffer (Ref. ^57^). *S. aureus*-Alexa Fluor 647 (100 μL, 5 mg/L) or anti-FitC-Alexa Fluor 647 (100 μL, 5 mg/L) was added and incubated for 30 min at room temperature followed by gentle washing with mM9 buffer. Unstained negative controls were incubated with mM9 buffer. Bacteria were visualized by confocal laser scanning microscopy (CLSM, Zeiss LSM700) equipped with a 63x/NA1.4 Plan-Apochromat objective using 488 and 639 nm excitation.

*S. aureus* biofilms were formed by inoculation of overnight culture into 96 well plates (IBIDI, 89626) for 2 h at 37° C. Non-adhered cells were then removed by rinsing, and wells were filled with BHI broth and incubated for 24 h at 37° C. Biofilms were then washed, blocked, and hybridized with antibodies, Biofilm embedded cells were stained with SYTO41 and visualized by CLSM using 405 and 639 nm excitation.

**Binding of ADC vs parent antibody to *S. aureus* detected by flow cytometry*.*** Single colonies of *S. aureus* from tryptic soy broth (TSB) agar plates were grown in TSB with 10 ug/mL of chloramphenicol for 19h at 37°C, 180 rpm. Bacteria were diluted to OD600=0.05 in TSB with chloramphenicol, and grown at room temperature to midlog phase (OD600=0.427 corresponding to 4.27*10^8^ CFU/mL) followed by wash and resuspension in RPMI-H medium (Roswell Park Memorial Institute medium (RPMI), added 0.05% HSA) and stored at -20°C. Freezer cultures were thawed at room temperature and diluted in RPMI-H to 3.75*10^7^ CFU/ml before the experiments.

*S. aureus* was added to 96-well plates (20 μL/well) and mixed with 20 μL of ADC or antibody (*S. aureus* polyclonal rabbit IgG, Thermo Fisher) in 3-fold dilution series for 30 min at 4°C, 600 rpm.. Wells were washed with 160 μL of RPMI-H at 2823 x g for 7 min. Goat-anti-rabbit-IgG-APC detection antibody (1 μg/mL ) (Molecular Probes, 199312) with 30 μL RPMI-H was added to the bacteria and incubated for 30 min at 4°C, 600 rpm. Unbound detection antibody was removed by washing and bacteria fixated with 100 μL of 1% paraformaldehyde (PFA) (Polysciences) in RPMI-H. Quantification of ADC and antibody bacterial binding was measured with flow cytometry (BD FACSVerse) and data analysed with FlowJo Software (Version 10.8.1).

**Phagocytosis of bacteria opsonized by ADC and parent antibodies.** Neutrophils where isolated from healthy blood donors using the Ficoll-Histopaque method. ^58^ For opsonisation, *S. aureus* was added to 96-well plates (20 μL /well) and incubated with 10 μL of 1% IgG/IgM depleted human pooled serum ^37^ and 10 μL ADC, specific or non-specific antibody in 3-fold dilution series for 15 min at 37°C, 750 rpm . 10 μL of neutrophils (7.5×10^6^ cells/ml) were added and incubated 15 min at 37°C, 750 rpm . Phagocytosis was stopped and cells fixated by adding 80 μL 1.62% cold PFA to each well. Quantification of phagocytosis was done using flow cytometry and data was analysed with FlowJo Software. The gating strategy used was based on neutrophil population gated on FSC-SSC and GeoMFI of neutrophils using the mAmetrine channel.

**Antimicrobial effect of ADC against planktonic *S. aureus*.** Overnight cultures were harvested by centrifugation (13150 x g, 8 min), resuspended in mM9 and diluted to OD_600_=0.005. 100 µL of each culture was added to Eppendorf tubes and centrifuged for 8 mins at 13150 x g. The formed pellets were resuspended in 100 µL of mM9. containing the chosen concentrations of specific ADC or non-specific ADC, with or without 5 mM NAC. In experiments with lower OD_600_ than 0,05 the pellet was not resuspended but ten μL of bacterial suspension was transferred to 90 μL containing the chosen concentration of *S. aureus* specific or non-specific ADC. Blanks (without bacteria) and growth controls (with bacteria) contained mM9 media. All samples were incubated for 2 h at 37°C, 180 rpm and centrifuged twice for 8 min, 1315 x g and resuspended in mM9. Ten-fold dilutions were made in mM9 of each sample, followed by plating 10 μL on agar and incubation for 19 h at 37 °C and then CFU enumeration. The antimicrobial effect was evaluated by comparing the CFU of treated samples with the CFU of the growth control from the same experiment.

**Antimicrobial efficacy of specific ADC against biofilm associated *S. aureus*.**Biofilms were grown on peglids by inoculating them in an overnight culture (OD_600_=1) for 1h at 37 °C,120 rpm. and then transferred to 96-well plates containing BHI and inoculated for48h at 37° C, 120 rpm. Media was exchanged after 24 h. The peg-lids were treated in 96-well plates with 2-fold dilutions of specific or non-specific ADC (0.5-2 mg/ml) and incubated 15 min at 37 °C. 120 rpm. Unbound ADC was removed by incubation in mM9 for 2 h. Peglids were transferred to a microtiter plate with mM9 and sonicated for 10 min to remove biofilm. Ten-fold dilutions were made from each treatment and spotted onto BHI agar plates followed by incubation at 37 °C before CFU enumeration was performed.

**Thiol labeling of planktonic *S. aureus*.** Overnight cultures were washed twice (13150 x g, 10 min) and resuspended in mM9 and diluted to OD_600_ = 1. Fluorescein maleimide (sample) or fluorescein (control) was added to the cultures for a final concentration of 0.1 mM. Bacteria where stained with SYTO60 (0.02 mM) and incubated for 30 min in the dark at 37 °C, 50 rpm. Stained bacteria were washed (14500 rpm, 10 min) and resuspended in PBS. 20 µL of sample and control was added to a superfrost ultra plus slide for 10 minutes. Bacteria were then visualised with CLSM through a 100× NA1.4 Zeiss Plan-Aprochromat oil objective, using 488 nm and 639 nm wavelengths for excitation and emission, respectively.

**Thiol labeling of *S. aureus* biofilm.** BHI supplemented with 5 % plasma (100 μL) was incubated in IBIDI 96 well black μ-Plate (IBIDI 89621) for 30 minutes. Wells were washed twice with fresh BHI and 200 μL of OD-adjusted *S. aureus* overnight culture (OD_600_=1) was added and incubated for 2 h at 37 °C. Media was replaced with fresh BHI and incubated for 24 h at 37 °C 50 rpm. BHI was removed and replaced with 1% blocking agent (BSA in mM9) and incubated 1h. Wells were washed three times in mM9, and 99 µL of mM9 was added. 1 μL fluorescein maleimide (0.1 mM) or fluorescein (0.1 mM) was added to the wells, and incubated 30 min in the dark at 37 °C, 50 rpm. Subsequently, wells were washed three times with PBS and resuspended in 1 x PBS (42 μL), and 8 μL SYTO60 (0.04 mM). Imaging was performed as described above.

***In vitro* toxicity test:**

The MOLT-4 cells, a human T lymphoblast cell line with acute lymphoblastic leukemia were grown as a suspension culture in pre-heated (37°C) RPMI-1640 Medium (Sigma R0883) containing 10% fetal bovine serum (Sigma F724), 2 mM L-glutamine (*Sigma G2150*) and 1% penicillin/streptomycin (*Sigma P0781*). The suspended cells were grown in 75 cm^2^ culture flasks and stored up-right in an incubator at 37 °C with 5% CO_2_.

**Cell counting.** Was carried out using a 1:1 ratio of Trypan Blue staining solution (Thermofisher) to cells and counting was fulfilled using an automated cell counter LUNA (Logos Biosystems*).* Cells were in same procedure controlled for viability, which was always higher than 80%.

**Toxicity test of ADC and mitomycin C.** MOLT-4 cells in passage P9-P11 (2.5x10^5^ cells/mL) were seeded in a 96-well plate followed by addition of a dilution series of either specific ADC or free mitomycin C (DMSO content were equalized). Control samples without specific ADC or free mitomycin C were also prepared with equivalent DMSO percentage. The samples were incubated with specific ADC or free mitomycin C for all 72 h at 37°C, 5% CO_2_. After this, cell viability was assessed using PrestoBlue cell viability reagent (Thermofisher A13262) using a BioTek Synergy H1 microplate reader (λex/ λem = 536nm/619 nm). The experiment was reproduced 3 independent times with 3 replicates each time.

***In vivo* experiments**

**Study animals.** Eight-ten week old C57bl/6j mice (Janvier Labs, Le Genest-Saint-Isle, France) were housed at The Animal Facilities Arhus University at standard room temperature, 12 h day/night cycle with water and food ad lib. The animals had an acclimatization period of one week prior to surgery. The study was approved by the Danish Animal Experiments Inspectorate under permission 2016-15-0201-01121 and done under supervison of the faculty vets.

**Inoculation of steel implants.** We used the isolate *Staphylococcus aureus* SAU060112, a clinical isolate from a patient with a prosthetic joint infection^31^. Cultures from tryptic soy broth (TSB) agar plates were grown overnight in TSB media for 18 h at 37°C, 180 rpm. The overnight culture was diluted in TSB to OD600=0.1 corresponding to 5×10^6^ CFU/mL, and was added to a 50 ml falcon tube with the steel implants (Ento Sphinx, Pardubice IV, Czech Republic) and grown for 18 h at 37°C. The implants were then transferred to fresh PBS before transportation to the animal facilities.

**Implant-associated osteomyelitis model.** The surgical procedure was based on a model by Jørgensen et. al ^29^. Briefly, mice were anesthetized with inhalation isoflourane (4-5% induction, 2% maintenance). Each animal was given an injection of buprenorphine 0.15 mg/kg s.c and the right hind leg was shaved. Implants were then surgically implanted transcortically through the left tibia at the proximal epiphysis. The implant was bent in a U shape and cut close to the skin on both sides. Afterwards, the skin and adjacent tissue was manipulated to cover the implant and animals where returned to their cages. Buprenorphine 0.009 mg/ml was administered in the water the first four days post-surgery.

**Deposition of antibodies at infection site.** Three days after infection animals were equally divided in to two groups by letter randomization. Group 1 (n=7 infected implant, n=3 sterile implant) was injected iv with antibody(*S. aureus* polyclonal rabbit IgG) conjugated to Alexaflour680. Group 2 (n=7 infected implant, n=3 sterile implant) was injected iv with unspecific antibody (FITC mice monoclonal IgG1, Thermo Fisher) conjugated to Alexaflour 680. The animals were scanned using *in vivo* imaging system (IVIS®, Xenogen, Alameda, CA, USA) at baseline and 1 h, 4 h and 24 h after injection. Fluorescent intensity from both legs of each animal was measured at excitation 675 nm, emission 720 nm. Analysis of data was done with Living Image Acquisition/Analysis Software Package. Fluorescence signal was counted by selecting the region of interest (ROI) of the implanted leg (ROI-1) and the non-implanted leg (ROI-2) of each animal. The Average radiant efficiency ([p/s/cm²/sr] / [µW/cm²]) of the ROI of each animal was then calculated for both legs and ROI-1 was subtracted from ROI-2 to remove background noise (Figure 5c).

**In vivo efficacy of antibacterial treatment.** Seven days after infection, 14 mice with an infected implant from the *in vivo* imaging were randomly divided into three treatment groups; 1) NaCl 0.9%, 0.5 mL 24 h/s.c (n=3), 2) Vancomycin (Bactocin®, MIP Pharma GmbH) 110 mg/kg/12h/s.c (n=5), 3) Vancomycin 110 mg/kg/12 h/s.c + specific ADC 5 mg/kg/24 h/iv (n=5). The final concentration of ADC was 5 mg/kg of which 4.916 mg/kg was antibody and 86 µg/kg was mitomycin C. Animals were treated for three days followed by a 36 h antibiotic washout period to avoid carry-over effect, and then euthanized by cervical dislocation. The tibia was exposed by surgical incision and the bone with implant was removed and snap-frozen at -80°C.

**Quantification of bacterial load.** Implants were carefully removed from the tibial bone and then submerged in 1 mL PBS (11 mM, pH 7.5), vortexed for 30 sec. and sonicated for 5 min at 45 kHz and 110 W (USCS1700 T, VWR). The tubes were vortexed 30 sec again and sonicate from each sample was serially diluted in triplicates, plated onto 5% blood agar plates and incubated at 37°C for 48 h. CFU was enumerated and bacterial load was calculated.

**Comparison of ADC with drug combination therapy.** Implants were inoculated as previously described, but with minor optimisations. Implants were inoculated in falcon tubes with five implants in each tube instead of all implants in one tube and then kept in the overnight culture until implantation instead of transferring them to PBS after inoculation. Animals were operated following Implant-associated osteomyelitis model. Seven days after infection, 40 animals were randomly divided into four treatment groups; 1) NaCl 0.9%, 0.5 mL 24 h/s.c (n=10), 2) Vancomycin 110 mg/kg/12 h/s.c (n=10), 3) specific ADC 5 mg/kg/24 h/iv (n=10), 4) Vancomycin 110 mg/kg/12 h/s.c + specific ADC 5 mg/kg/24 h/iv (n=10). Following 3 days of treatment and 36h of antibiotic washout, animals were then euthanized and bone with implant was removed and bacterial load was analysed immediately after, following the quantification of bacterial load protocol.

**Quantification of treatment efficacy of ADC components.** The experiment was carried out similar to the Comparison of ADC with drug combination therapy study, with the same optimisations but with the following treatment groups: 1) NaCl 0.9%, 0.5 mL 24 h/s.c (n=10), 2) specific ADC 5 mg/kg/24 h/iv (n=10), 3) mitomycin C 86 µg/kg/24 h/iv (n=10), 4) specific antibody 5 mg/kg/24 h (n=10), 5 non-specific ADC 5 mg/kg/24h (n=10).

**Statistical analysis.** All viability data was prepared by subtracting background from the raw data and normalizing to the independent viability controls. The viability was then plotted as a function of the logarithm of the concentration. The independent IC50 values were estimated by fitting to a sigmoidal curve (four parameters, variable slope) using the software GraphPad Prism 9.

**NMR spectra**

**NMR of 1b**

**^1^H-NMR:**

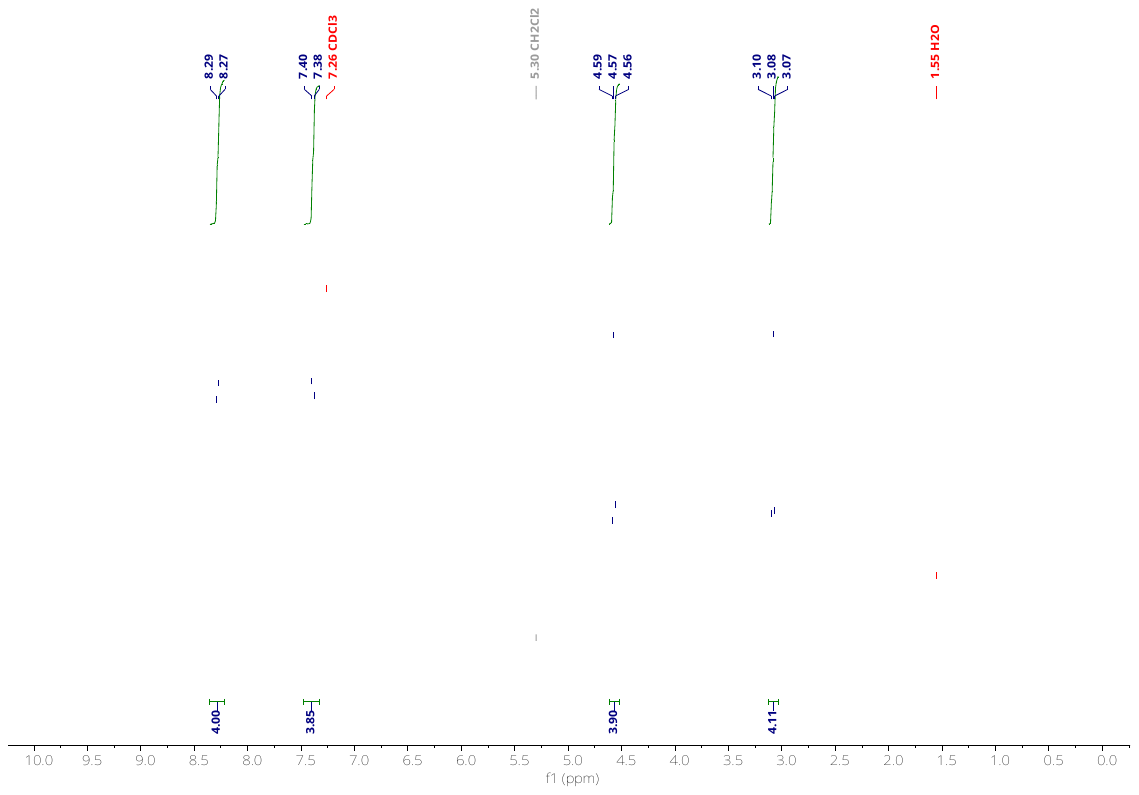

**^13^C-NMR:**

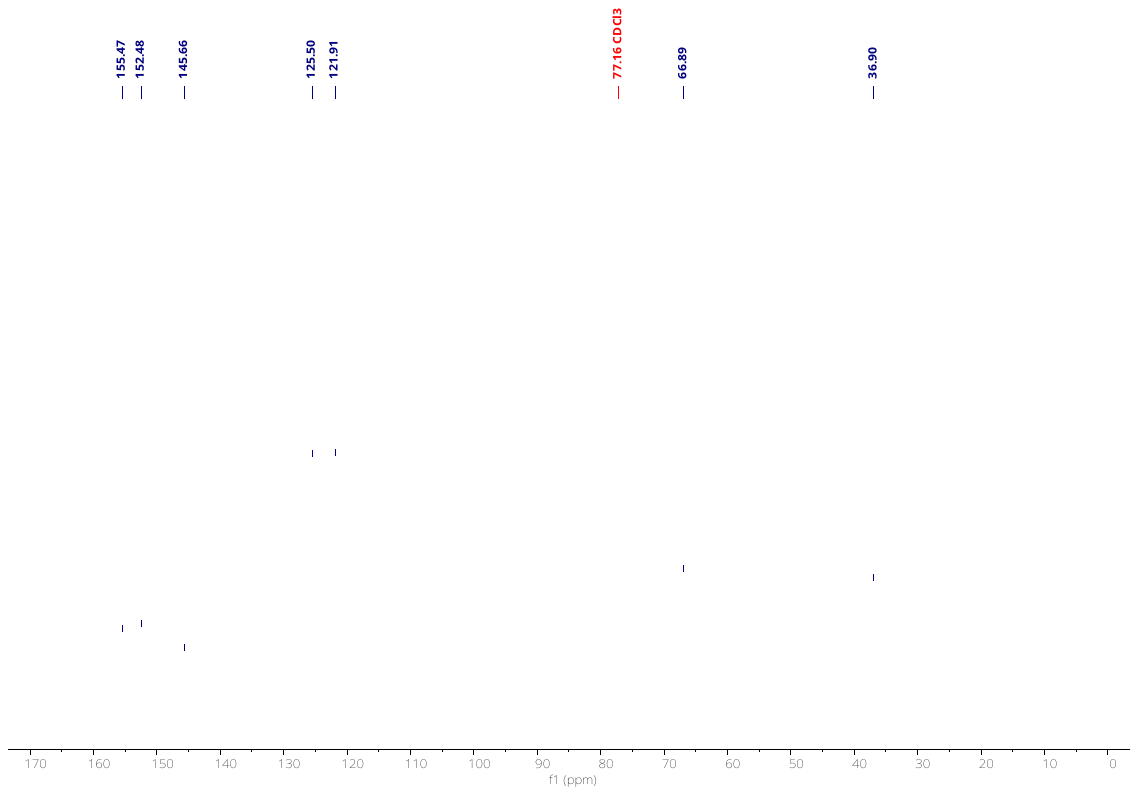

**NMR of 1**

**^1^H-NMR:**

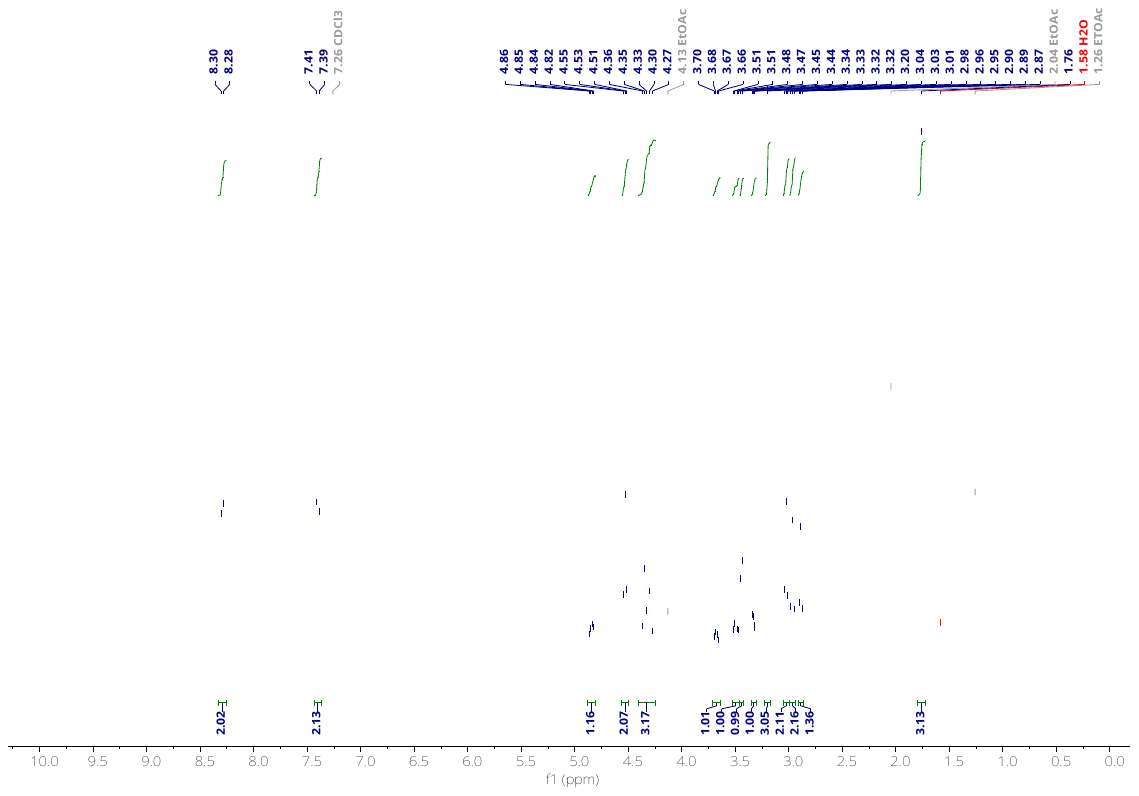

**^13^C-NMR:**

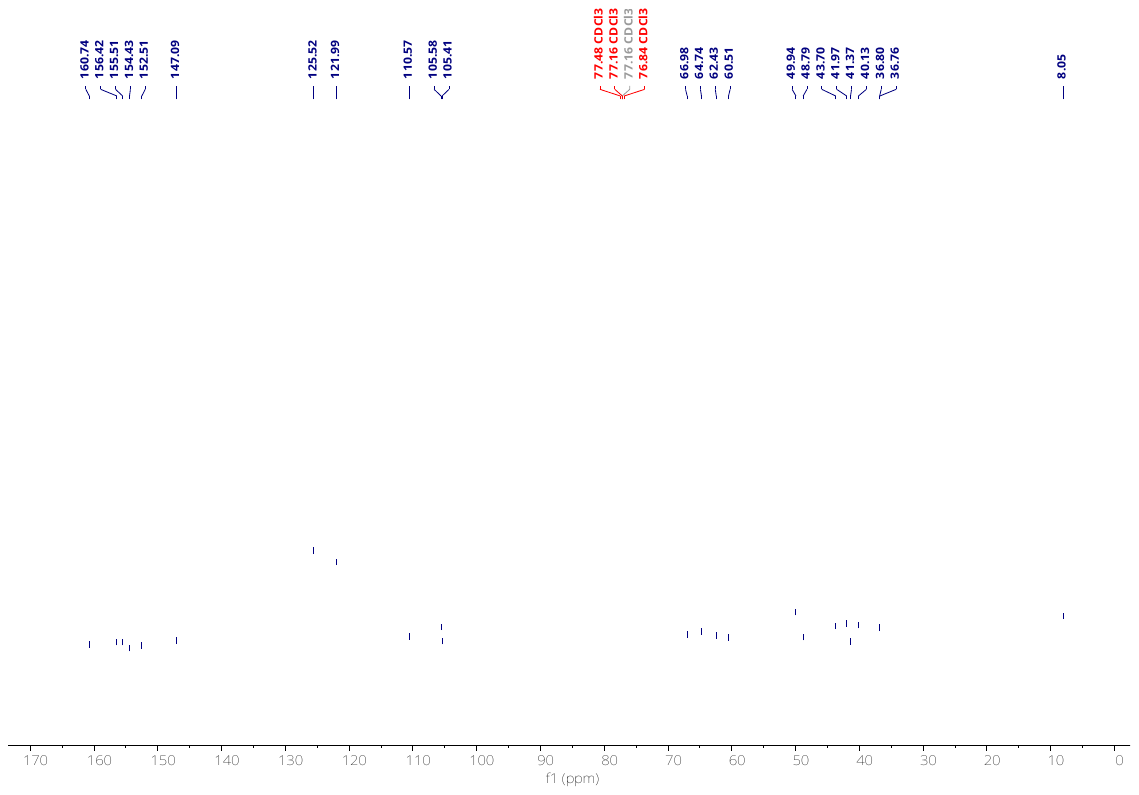

**NMR of 2b**

**^1^H-NMR:**

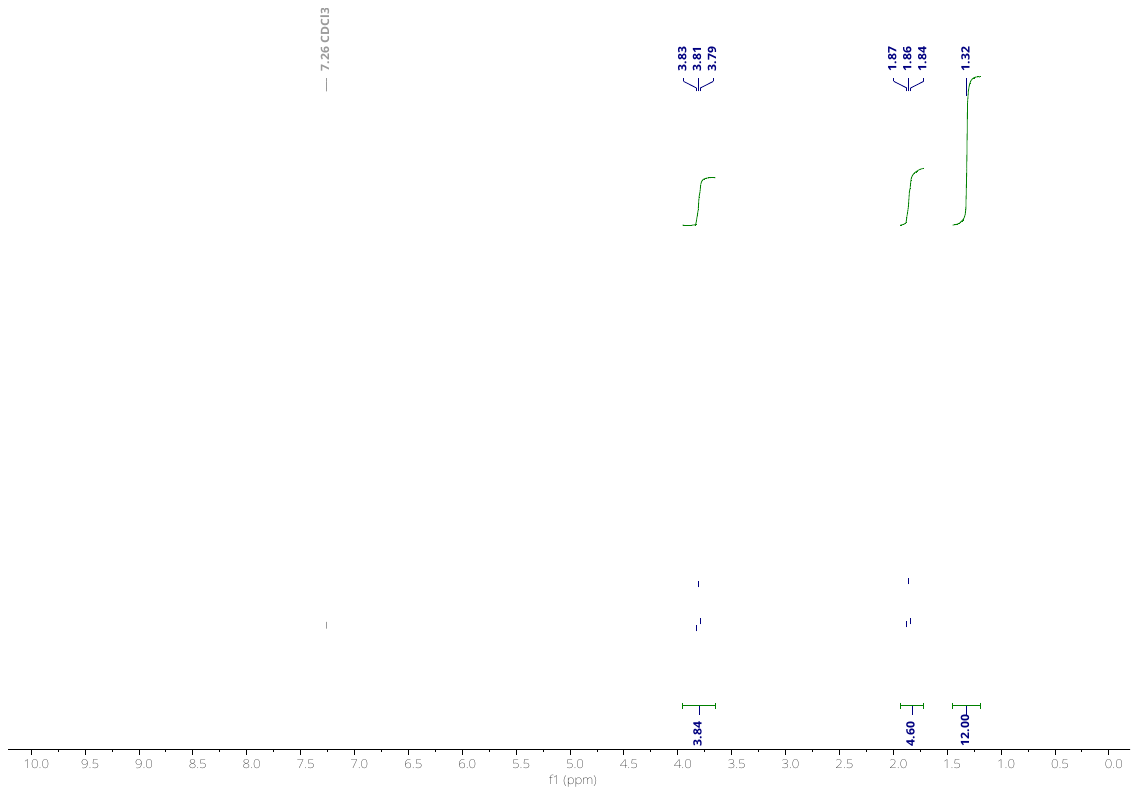

**^13^C-NMR:**

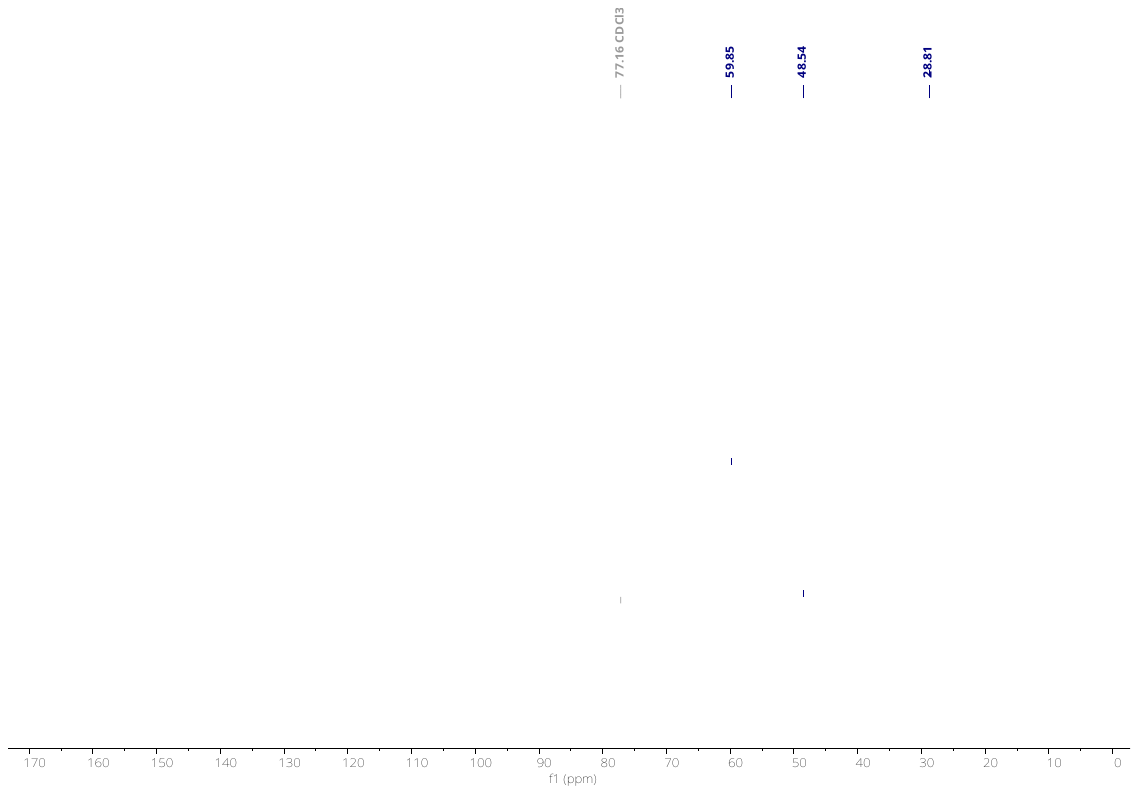

**NMR of 2c**

**^1^H-NMR:**

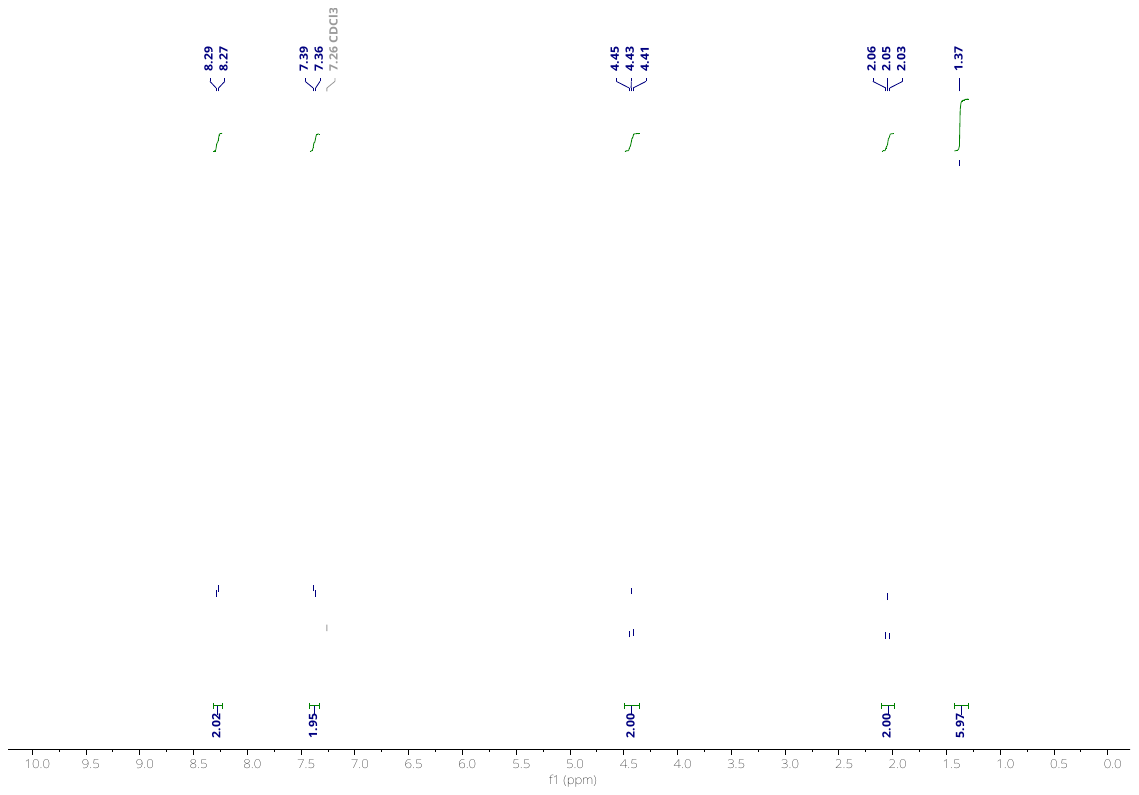

**^13^C-NMR:**

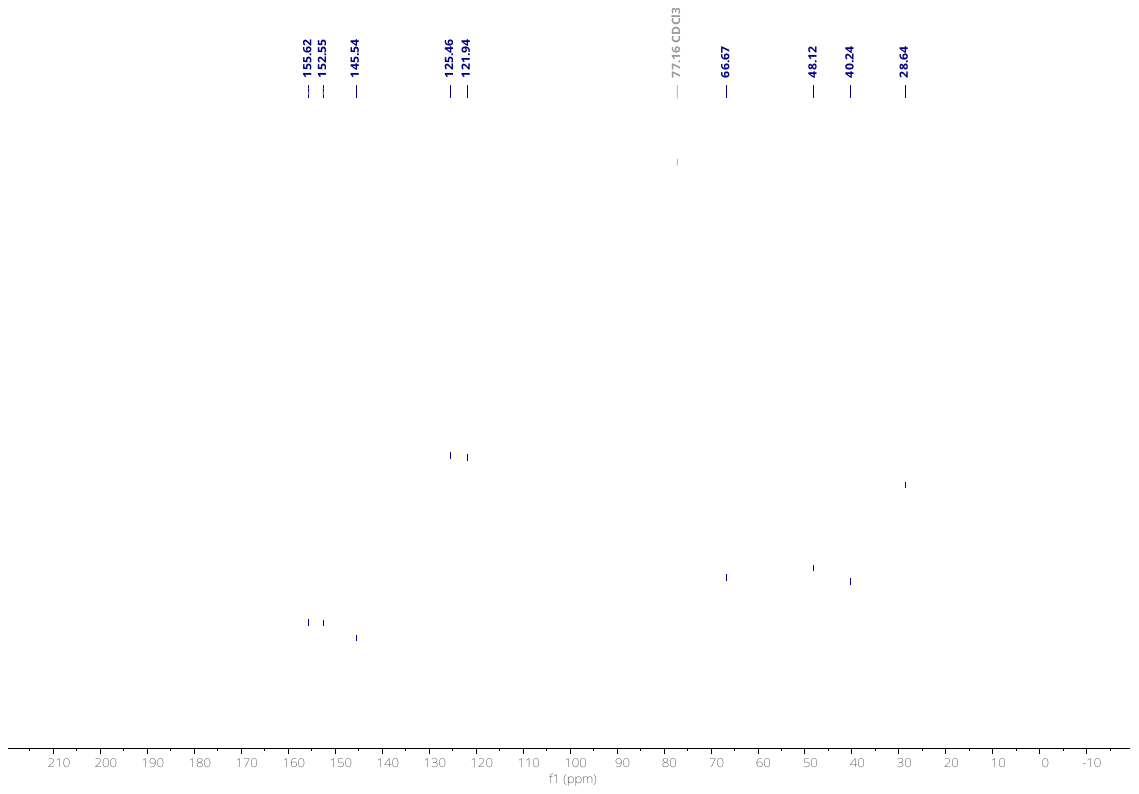

**NMR of 2**

**^1^H-NMR:**

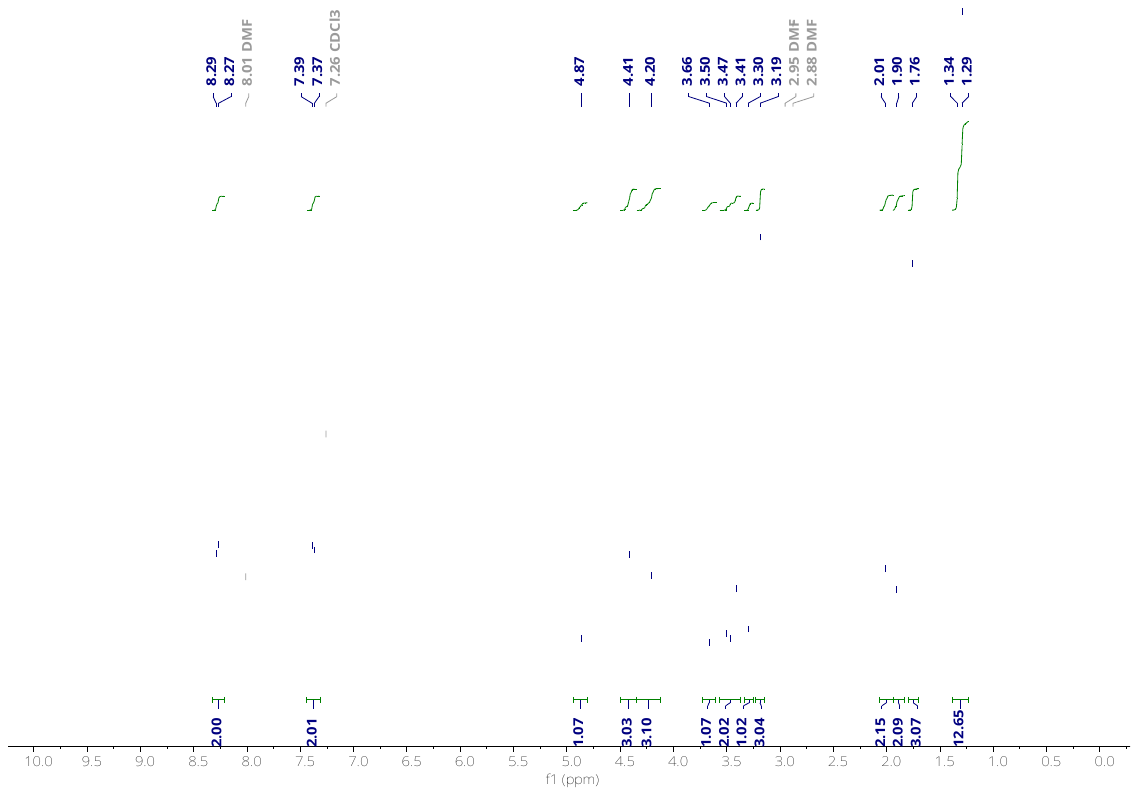

**^13^C-NMR:**

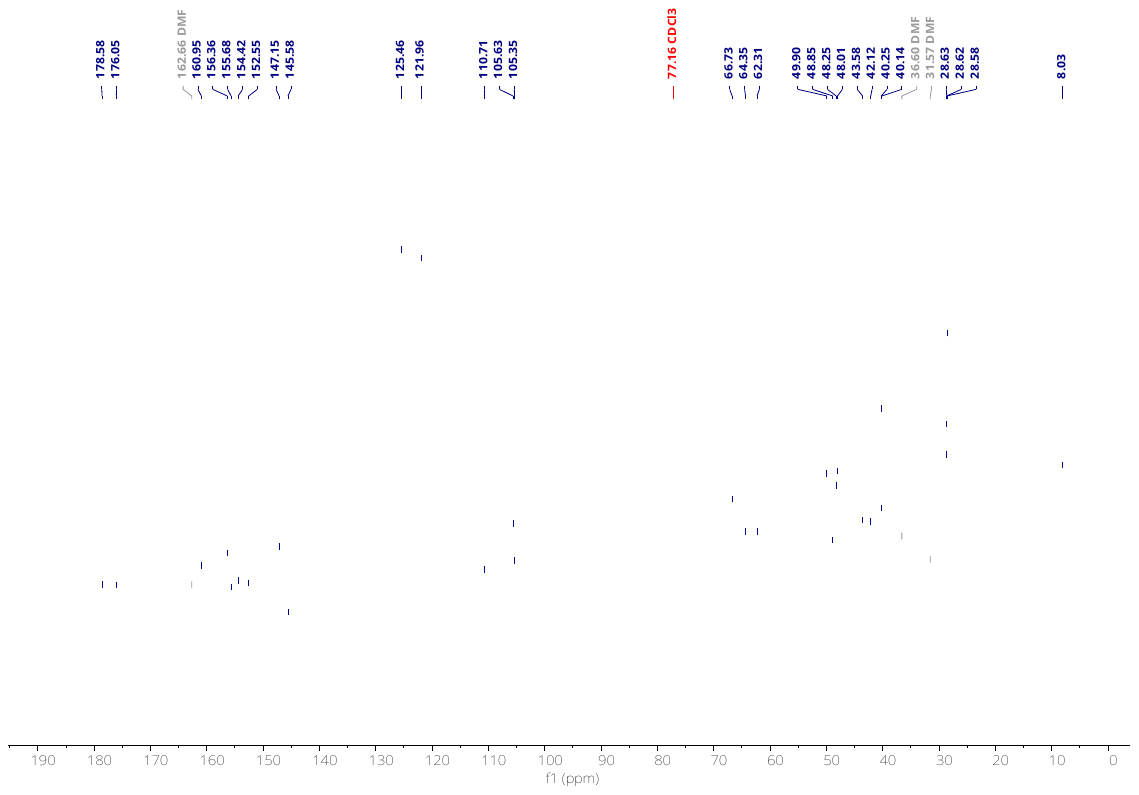
